# A Diploid *Panax* Genome Reveals Ginsenoside Diversity Driven by UGT Family Diversification and Network Rewiring Rather than Gene Family Expansion

**DOI:** 10.64898/2026.06.16.732563

**Authors:** Zhe Xu, Wei Li, Fu-gang Wei, Gao Xiong, Zhong-jian Chen, Li-zhi Gao

## Abstract

The medicinal herb *Panax notoginseng* produces a structurally diverse array of triterpene saponins (ginsenosides), yet the genetic basis of this chemical complexity remains unclear. Here we present a high-quality chromosome-level genome of diploid *P. notoginseng* and integrate comparative genomics with multi-tissue, multi-year metabolomics and transcriptomics. Surprisingly, unlike tetraploid *Panax* species, *P. notoginseng* shows no general expansion of core saponin biosynthetic gene families. Instead, lineage-specific diversification of UDP-glycosyltransferase (UGT) families, a recent burst of LTR retrotransposons, and enrichment of species-specific genes in metabolic modification pathways point to an alternative evolutionary route. Saponin accumulation follows strict spatiotemporal compartmentalisation, and co-expression network analysis reveals that the biosynthetic machinery is not static but continuously rewired during development-from a basic synthesis module in the first year to a modular pattern supporting both broad accumulation and branch-specific modification by the third year. Seventeen differentially expressed *UGTs* show clear tissue preferences and saponin-branch correlations. As a representative example, *PnUGT33* is tightly linked to the PPD-type saponin branch; structural modelling, molecular docking and 100 ns molecular dynamics simulations demonstrate its differential recognition of diverse triterpene skeletons. Collectively, our findings establish that ginsenoside diversity in diploid *P. notoginseng* arises primarily from *UGT* lineage diversification, developmentally rewired regulatory networks and *UGT* mediated branch selective post-modification, rather than from expansion of core pathway genes. This work provides a new paradigm for understanding how plants achieve metabolic complexity without whole genome duplication or massive gene amplification.

## Introduction

The genus *Panax* (Araliaceae) is one of the most medicinally valuable plant groups; its major active constituents are triterpene saponins (ginsenosides), which exhibit a broad range of pharmacological activities including anti-inflammatory, anti-tumour, cardioprotective and immunomodulatory effects^1–7^. The structural diversity of ginsenosides is the chemical basis for their multiple pharmacological functions. Based on the aglycone skeleton, ginsenosides are classified into dammarane-type: protopanaxadiol (PPD) and protopanaxatriol (PPT) types, and oleanane type; each class comprises dozens of molecular entities that differ in the number, type and linkage positions of sugar moieties^1,2^. Nevertheless, the genetic basis and mechanisms underlying this chemical diversity remain largely unknown.

*Panax notoginseng* (Sanqi) is a representative medicinal species of the genus *Panax*. Its dried roots are a key ingredient in traditional Chinese medicine preparations such as Yunnan Baiyao and Sanqi Shangyao Pian^7, 8^ .Compared with closely related species such as *P. ginseng* and *P. quinquefolius*, *P. notoginseng* exhibits a distinctive ginsenoside composition profile. It not only accumulates the species-specific notoginsenoside R1 (NR1) but also displays marked species specificity in both the PPD/PPT-type ginsenoside ratio and glycosylation patterns^9–12^. This specificity is evident not only at the interspecific level but also across different tissues and developmental stages within the same plant. The underground tissues of P. notoginseng (main root, fibrous root and rhizome) predominantly accumulate PPD and PPT ginsenosides, whereas the aerial tissues are dominated by PPD-type ginsenosides. Furthermore, total ginsenoside content exhibits dynamic variation with growth years; levels in second year and third year plants fluctuate relative to each other across different harvest months, and the component ratios are continuously remodeled throughout the growth process^9,11^. This complex spatiotemporal accumulation pattern suggests that ginsenoside biosynthesis in *P. notoginseng* is highly regulated, but the genetic basis and molecular mechanisms remain unclear.

In recent years, several genome assemblies of *P. notoginseng* have been published, providing important resources for dissecting ginsenoside biosynthesis^2,13,14^. Zhang et al. (2017) reported the first draft genome, revealing a genus-specific whole-genome duplication event^2^. Jiang et al. (2020) and Yang et al. (2021) further provided chromosome-level reference genomes^13,14^. However, these genomic resources have mostly been used for functional characterisation of individual structural genes or for expression analysis in limited tissues^13,15,16^, and a systematic understanding of how the species-specific and spatiotemporally distinct saponin accumulation pattern arose is still lacking.

A general framework of ginsenoside biosynthesis has been established. Isoprenoid precursors are generated via the mevalonate (MVA) and 2-C-methyl-D-erythritol-4-phosphate (MEP) pathways, followed by triterpene skeleton cyclisation, oxidative modifications by cytochrome P450s (CYPs), and finally glycosylation by UDP-glycosyltransferases (UGTs), which produce structurally diverse saponins^17,18^. The terminal glycosylation step determines the sugar chain composition, linkage pattern and final structure, and is considered a key driver of saponin chemical diversification^19,20^. In *P. notoginseng*, synthetic biology platforms have been used to elucidate the complete biosynthetic pathways of major saponin glycosylation products, and a UDP-xylose-dependent glycosyltransferase involved in notoginsenoside R1 biosynthesis has been identified^16^. These studies indicate that *UGTs* are not merely terminal modifying enzymes but are key nodes linking metabolic pathway output to chemical diversity.

Despite this progress, three critical gaps remain in our understanding of the mechanisms that generate saponin diversity in *P. notoginseng*. First, at the evolutionary level, genes involved in triterpene metabolism may be influenced by gene duplication and family expansion, but the trajectories differ among *Panax* species^21–24^. Comparative genomics analyses have revealed independent evolutionary lineages of key enzyme genes in *Panax* species, but it is unclear whether the unique saponin profile of the diploid *P. notoginseng* is driven mainly by expansion of core structural gene families or rather by lineage-specific diversification and functional reprogramming of downstream UGT families. Second, at the regulatory level, transcriptomic studies have shown tissue-specific expression patterns of saponin biosynthetic genes^5^, but whether UGT family members exhibit spatiotemporal expression patterns that mirror the metabolic landscape and how these patterns relate to different saponin branches remain unknown. Third, at the functional level, only a minority of the predicted *UGTs* in *P. notoginseng* have been functionally validated; the substrate scope, branch preferences and relationship to saponin structural diversity have only been sporadically reported^16,25^.

To address these gaps, we generated a high-quality chromosome-level reference genome of *P. notoginseng* and combined targeted metabolomics and transcriptomics of multiple tissues across three growth years to systematically dissect the genetic basis of saponin diversity. Focusing on the central role of *UGTs*, we aimed to understand the mechanisms of modifier-driven metabolic diversification from three perspectives: family evolution, spatiotemporal regulation, branch selective post-modification.

## Results

### Ginsenoside diversity in diploid *P. notoginseng* does not depend on expansion of core pathway gene families

To explore the genetic basis for the chemical diversity of saponins in P*. notoginseng*, we first assembled a chromosome-level reference genome of approximately 2.27 Gb, with >98% of the sequence anchored to 12 chromosomes (Fig. 1a, Supplementary Tables S1-S3, Supplementary Fig. S1, S2). BUSCO completeness was 92.1% (Supplementary Table S4). High mapping rates of Illumina short reads (∼99.12%) and full-length transcripts (∼85.49%) to the assembly supported its high quality (Supplementary Table S5). Annotation yielded 46,425 protein-coding genes, as well as 1,800 non-coding RNAs (Supplementary Table S6). Comparative genomics analyses suggested that the genus Panax originated 52-55 million years ago, with the closest evolutionary relationship being between *P. notoginseng* and *P. vietnamensis* var. *fuscidiscus*, from which it diverged approximately 29.91 million years ago (Fig. 1b). A clear 1:2 syntenic relationship was observed between *P. notoginseng* and three tetraploid *Panax* species (Fig. 1c). Repetitive sequences accounted for 87.51% of the genome, with Ty3-gypsy LTR retrotransposons comprising 60.22% and having undergone a marked expansion approximately 0.10-0.23 million years ago (Supplementary Fig. S3, Supplementary Table S7), indicating a lineage-specific recent transposon burst.

**Fig. 1.**
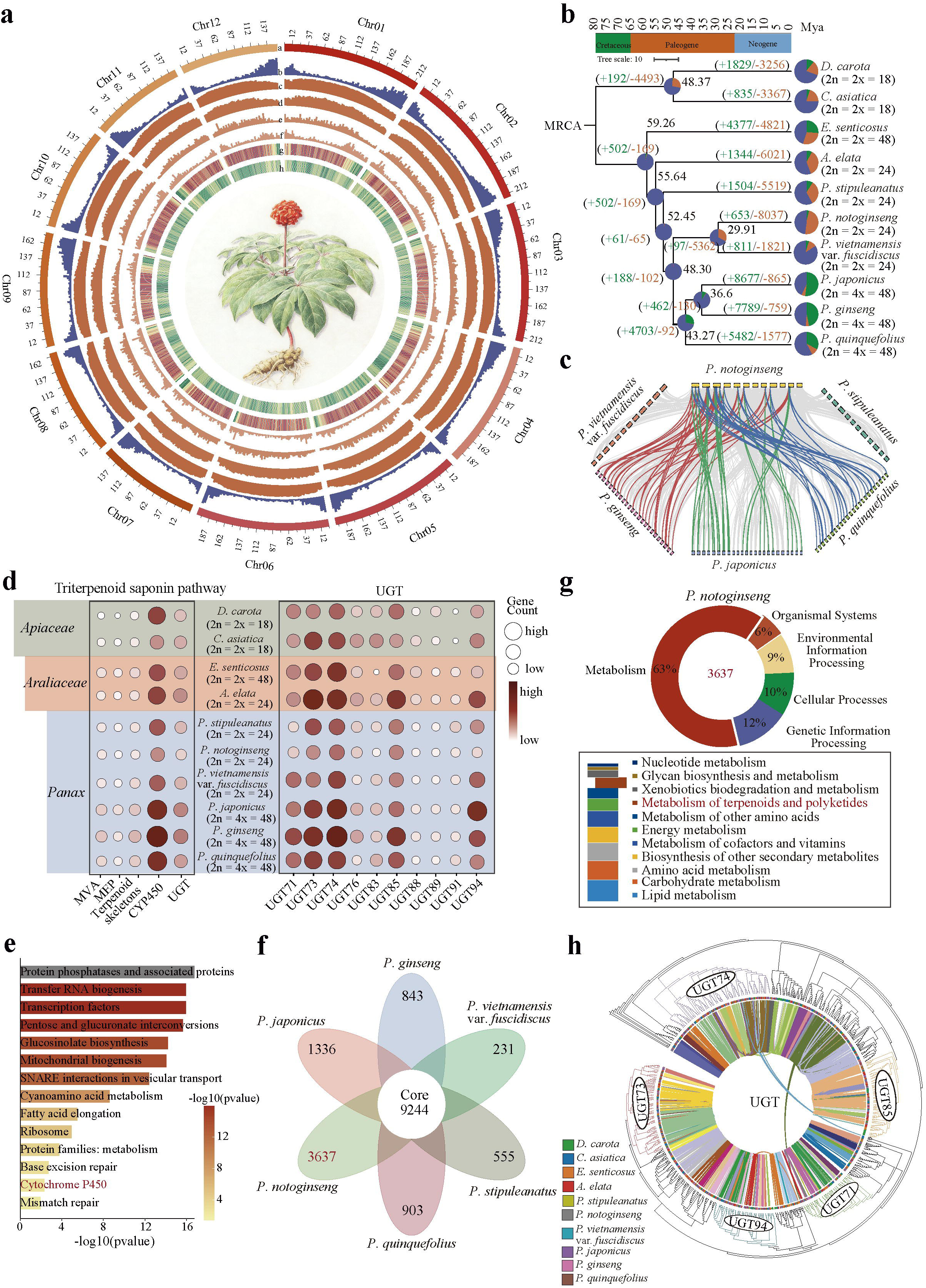
Genome features of *P. notoginseng* and evolution of the triterpene saponin biosynthesis pathway. **(a)** Pseudo-chromosome map of *P. notoginseng*. a, Circular representation of 12 pseudo-chromosomes; b, distribution of protein-coding genes; c, distribution of transposable elements (TEs); d, distribution of Ty3-gypsy-like LTR retrotransposons; e, distribution of Ty1-copia-like LTR retrotransposons; f, distribution of DNA transposons; g, number of SSRs per 3,000-kb window; h, GC content. **(b)** Phylogenetic tree of ten plant species. Gene family expansions and contractions are indicated in red and green, respectively. Numbers on the nodes represent the divergence time (million years ago, MYA). **(c)** Syntenic relationships among *Panax* species chromosomes. Curves indicate syntenic blocks, and highlighted curves denote 1:2 syntenic relationships between *P. notoginseng* and polyploid species. **(d)** Copy number comparison of ginsenoside biosynthesis-related genes across ten plant species. Larger and darker bubbles indicate higher copy numbers. **(e)** Functional enrichment analysis of expanded gene families in *Panax* species. The colour scale indicates the significance of enrichment, measured by the magnitude of -log10(q-value). **(f)** Venn diagram of orthologous gene distribution among *Panax* species. Overlapping numbers indicate shared orthogroups. **(g)** Functional annotation of *P. notoginseng*-specific gene families. The upper circular plot shows enriched pathways by KEGG first level categories, with different colours representing distinct pathway categories; the lower bar chart shows a detailed view of second level categories within metabolic pathways. **(h)** Phylogenetic relationships and syntenic features of the UGT gene family. Branches in different colours represent distinct UGT subfamilies. Connections within the circle indicate syntenic gene pairs, with colours corresponding to different groups defined by algorithmic clustering.

We found that core pathway gene families for ginsenoside biosynthesis have not expanded in *P. notoginseng*. We assessed copy numbers of 20 core saponin pathway gene families across ten species, including *Panax* and outgroups from Apiaceae. In marked contrast to tetraploid *Panax* species, the diploid *P. notoginseng* did not show significant expansion in most of these families (Fig. 1d). Several UGT subfamilies (e.g., UGT73, UGT83, UGT93 and UGT94) even showed contraction (Fig. 1d, Supplementary Table S8). Although expanded gene families in Panax were enriched in pathways related to secondary metabolism such as cytochrome P450 and carbohydrate metabolism and transcriptional regulation (Fig. 1e), *P. notoginseng* possessed the largest number of species-specific gene families (3,637) among the six *Panax* species examined (Fig. 1f, Supplementary Table S9). These families were significantly enriched in terpenoid metabolism, cytochrome P450-mediated xenobiotic metabolism and glycan biosynthesis (Fig. 1g, Supplementary Table S10), suggesting that the genetic innovation in *P. notoginseng* is more focused on the modification layer rather than on expanding the copy numbers of core-skeleton genes.

We also observed lineage-specific differentiation of the UGT family in *P. notoginseng*. In total, 64 *UGT* genes were identified; phylogenetic analysis showed that the newly identified *PnUGTs* are not confined to a single clade but are distributed across several UGT lineages, forming multiple lineage-specific branches (Supplementary Fig. S4). Phylogenetic and synteny network analyses further revealed a lack of significant syntenic relationships among different UGT subfamilies (Fig. 1h, Supplementary Table S11), suggesting that the UGT family may have undergone independent diversification events rather than simple conservative retention or global expansion. Together, these results indicate that the saponin diversity of *P. notoginseng* is unlikely to be attributable to expansion of core biosynthetic gene families, but may be more closely linked to the lineage-specific diversification of modifier enzymes such as *UGTs*.

### Strict spatiotemporal compartmentalization of ginsenoside accumulation and its shaping by dynamic co-expression network rewiring

Targeted metabolomics of nine tissues (main root, fibrous root, rhizome, leaf, stem, flower, pedicel, berry, seed) from first, second and third year (year 1-3) plants quantified 18 signature ginsenosides. Underground tissues mainly accumulated PPD-type ginsenosides (Rd, Rb1, Ra3), PPT-type ginsenosides (Re, Rh1, Rg2) and the *P. notoginseng* specific NR1, which together accounted for on average >97.1% of total underground saponin content. Aerial tissues mainly accumulated PPD-type saponins (Rd, Rb1, Ra3, Rb3, Rc, Rb2), which accounted for >91.8% of total aerial saponin content (Fig. 2a, Supplementary Table S12). Sample correlation heatmaps showed that metabolic profiles clustered clearly by tissue type, with underground tissues, aerial vegetative tissues and reproductive tissues (flower, pedicel, berry, seed) separated from each other (Fig. 2b). Total saponin content peaked in the second year in underground and aerial vegetative tissues, and in the third year in reproductive tissues (Supplementary Table S12). Principal component analysis and correlation analysis further revealed highly coordinated accumulation of the PPD-type saponins Rb2, Rb3 and Rc (r = 0.99-1.00) (Fig. 2d, e). Thus, saponin accumulation in *P. notoginseng* is strictly tissue-compartmentalised and developmentally stage-dependent.

**Fig. 2.**
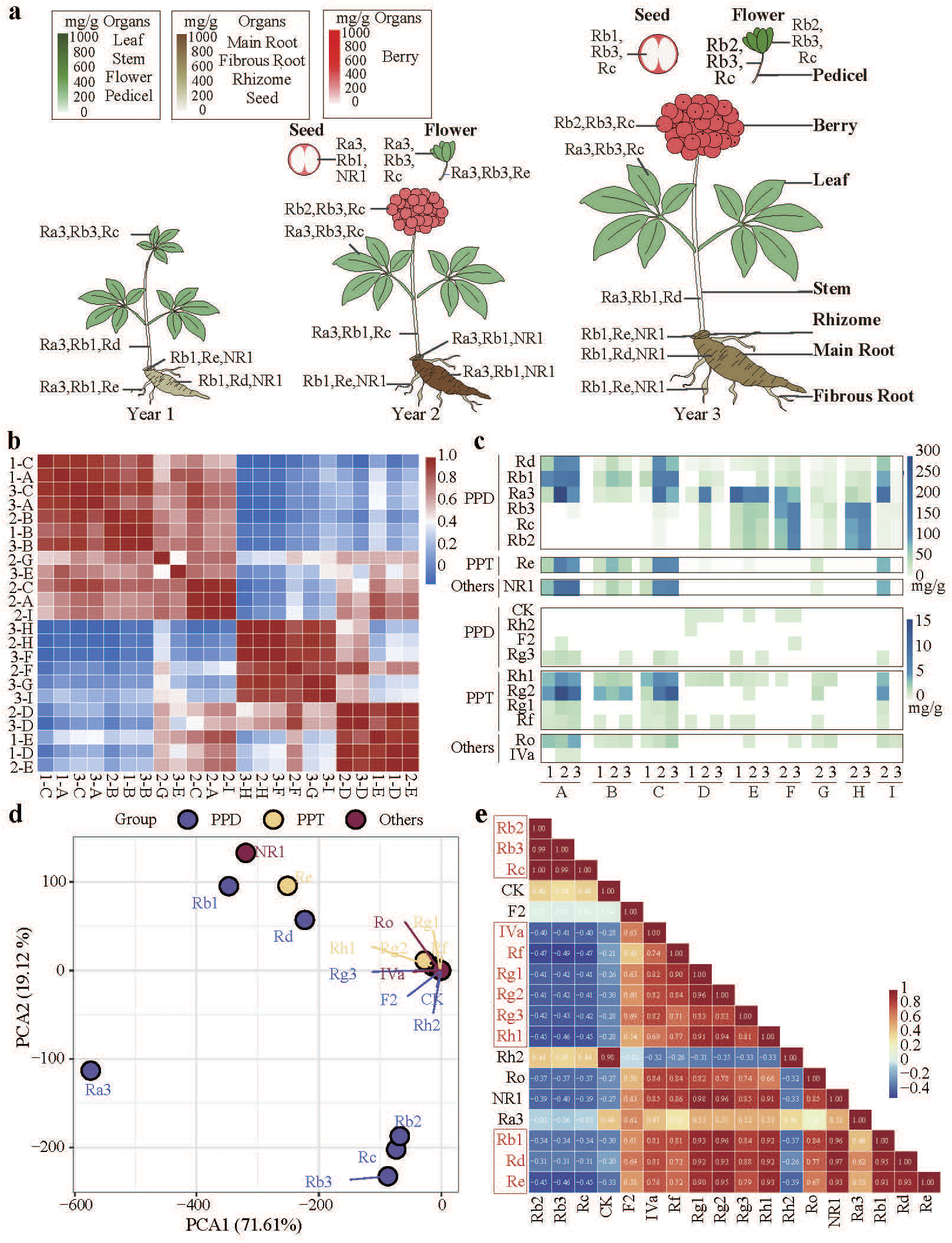
Spatiotemporal accumulation characteristics of triterpenoid saponins in *P. notoginseng*. Years 1-3 indicate the growth years. A-I represent: main root, fibrous root, rhizome, leaf, stem, flower, pedicel, berry, seed. **(a)** Heatmap of total saponin content in nine tissues of 1-3 year-old plants. Colour intensity corresponds to total saponin content. **(b)** Correlation heatmap among metabolome samples. Colour gradient indicates Pearson’s correlation coefficients. **(c)** Saponin composition and distribution profiles across different tissues and developmental stages. Samples are arranged by tissue type and stage. Saponins were divided into high-content (>15 mg/g) and low-content groups. **(d)** Principal component analysis (PCA) based on saponin distribution. Different colours represent different types of ginsenosides. **(e)** Correlation analysis between the contents of various saponins. Values indicate correlation coefficients, colour gradient reflects strength of correlation; highly correlated groups are highlighted in red.

Transcriptomic analysis of 59 samples (Supplementary Table S13) showed that sample variation was primarily driven by tissue type, with growth year having a weaker effect (Fig. 3a, b). qRT-PCR validated the transcriptome data (Supplementary Fig. S5, Supplementary Table S14). K-means clustering identified one expression cluster (Cluster 1) whose overall expression increased with growth year; this cluster could be divided into three functional sub-modules: redox and metabolic skeleton modification (Sub-cluster A), upstream signal transduction (Sub-cluster B) and glycosyltransferase-related functions (Sub-cluster C), the latter containing 22 glycosyltransferase-related genes (Fig. 3c, d). Homology searches using known saponin biosynthetic genes (Supplementary Table S15) and K-means clustering revealed that core pathway genes were not concentrated in a single cluster (Fig. 3c, f), but were dispersed across multiple modules, showing pronounced tissue and developmental stage specificity; among them, UGT family members exhibited the most divergent expression patterns (Fig. 3e, f, Supplementary Table S15). To further dissect this regulatory landscape, we constructed WGCNA co-expression networks for the first, second and third year plants using 11,674 genes each. The analyses identified 10, 14 and 21 co-expression modules at the three stages, respectively. And each showed significant differential correlations with saponin components and tissue types, indicating that saponin metabolism in *P. notoginseng* is not controlled by a fixed set of genes but by a multi-module network that is continuously rewired during development (Supplementary Fig. S6-S8).

**Fig. 3.**
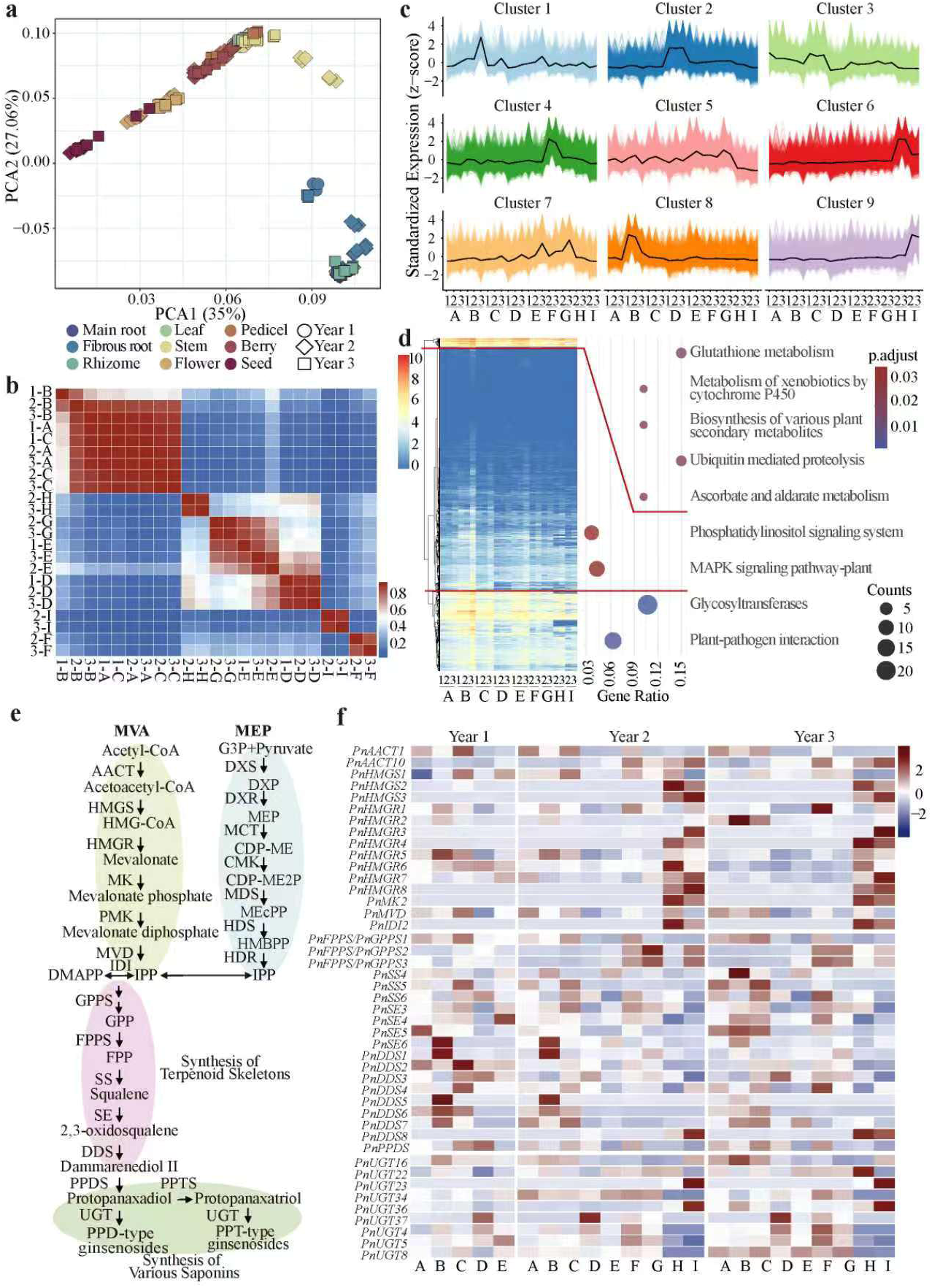
Transcriptome landscape of *P. notoginseng* and elucidation of the triterpene saponin biosynthesis pathway. Years 1-3 indicate growth years. A-I represent the same tissues as in Fig. 2. **(a)** PCA of transcriptome samples. Different developmental years and tissue types are distinguished by colours and shapes, respectively. **(b)** Correlation heatmap among transcriptome samples. Colour gradient indicates Pearson’s correlation coefficients. **(c)** Gene expression patterns based on K-means clustering (nine clusters). Horizontal axis: tissue samples across developmental stages; vertical axis: normalised expression values. **(d)** Functional subclassification of Cluster 1 from (c). Genes are divided into three categories based on expression patterns. **(e)** Schematic diagram of the triterpene saponin biosynthesis pathway in *P. notoginseng*. **(f)** Expression profiles of ginsenoside biosynthesis genes across different developmental years and tissues. Colour scale indicates gene expression levels.

In first year plants, 10 modules were identified. Y1-MEblue was positively correlated with Rc, Rb2, CK, Rh2, Ra3 and Rb3 and was associated with the rhizome tissue, representing the predominant ginsenoside-related module (Fig. 4a, Supplementary Fig. S6). This module integrated *PnAACT*, *PnMVD*, *PnMDS1*, *PnSS5* and multiple *CYPs*/*UGTs*, and was enriched in starch and sucrose metabolism and glycolysis, constituting a fundamental precursor supply, backbone biosynthesis, initial post-modification unit (Fig. 4b; Supplementary Tables S16, S17). At the second year, 14 modules were constructed. Y2-MEblue became the core productive module, positively correlated with Rb1 and Rh1. In leaf tissue, it integrated the precursor-supply genes *PnHDS2*/*3* and *PnFPPS*/*PnGGPS6*, the skeleton-formation genes *PnOSC3*/*9*, *PnSE2*, *PnSS1/2*, and numerous *CYPs* and *UGTs.* This formed a continuous unit for precursor supply, backbone formation and oxidative/glycosylation modification (Fig. 4a, Supplementary Fig. S7; Supplementary Table S16). Meanwhile, Y2-MEred was highly correlated with Rg1, NR1, Rf, Re, Rb1, Ro, IVa, Rg2, Ra3, Rd and flower tissue (Fig. 4a, Supplementary Fig. S7; Supplementary Table S16). KEGG enrichment showed that Y2-MEblue was enriched in source processes such as photosynthesis and carbon fixation, whereas Y2-MEred was enriched in high-energy pathways such as glycolysis, oxidative phosphorylation and the TCA cycle (Fig. 4c, Supplementary Table S17). In the Y2-MEblue module, pathway genes formed a large co-expression network with transcription factors including AP2/ERF, MYB, NAC, WRKY and bHLH (Fig. 4e).

**Fig. 4.**
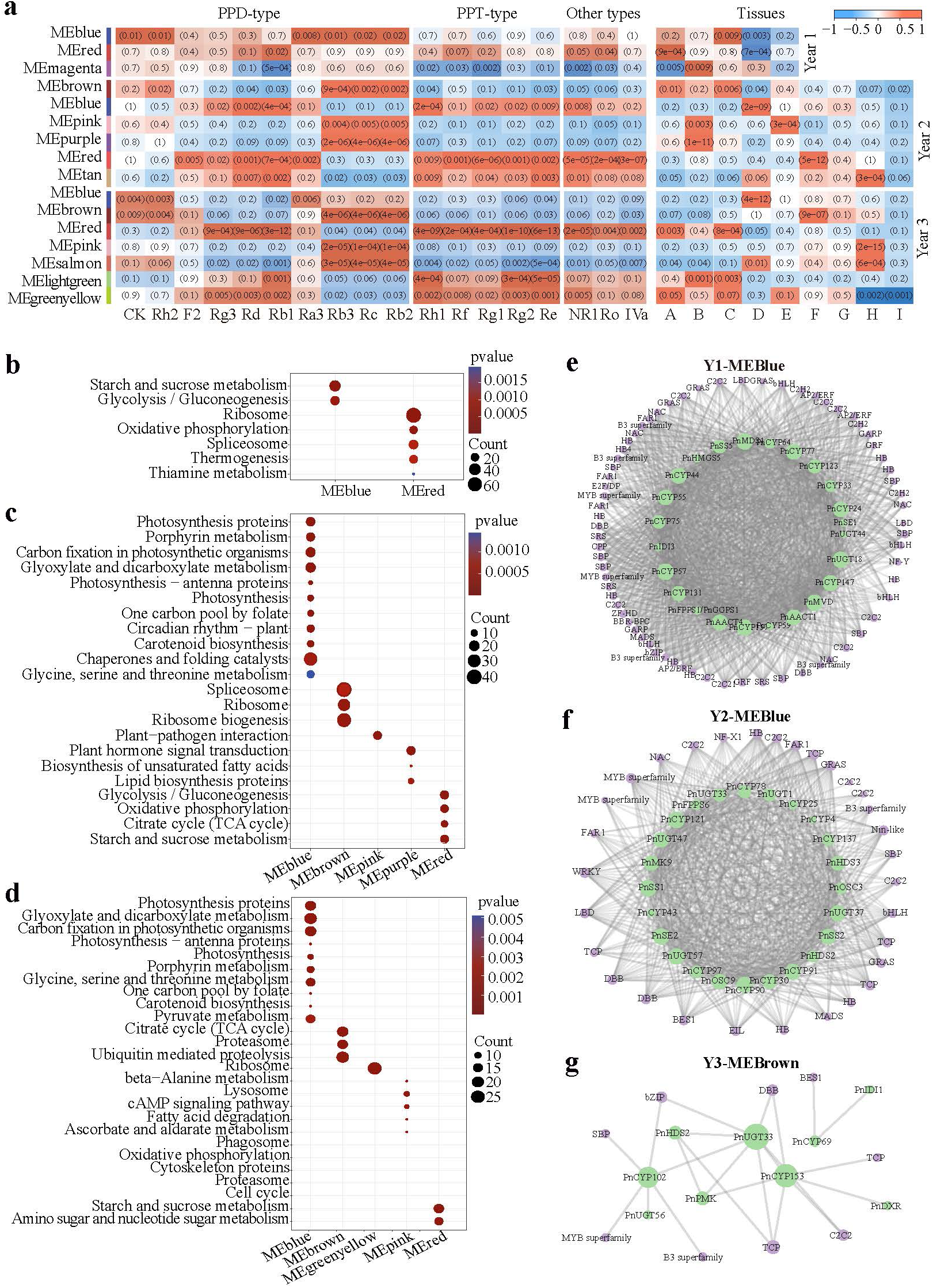
Identification and functional analysis of transcriptional regulatory modules associated with saponin accumulation in *P. notoginseng*. Years 1-3 indicate growth years; A-I as in Fig. 2. **(a)** Co-expression modules significantly correlated with saponin content and tissue type across different growth years. Values represent p-values; colour scale: red = positive correlation, blue = negative correlation. **(b–d)** Functional enrichment of key co-expression modules in year 1 (b), year 2 (c) and year 3 (d). **(e–g)** Interaction networks between pathway genes and transcription factors in year 1 (e), year 2 (f) and year 3 (g) samples. Transcription factors are indicated as purple nodes; node size represents connectivity.

By the third year, the metabolic network had further differentiated into a mature pattern comprising a “broad-accumulation module” and a “branch-modification module”. Y3-MEred, Y3-MElightgreen and Y3-MEgreenyellow were positively correlated with Rg1, NR1, Rf, Re, Rg3, Rb1, Rh1, Ro, IVa, Rg2 and Rd, and were significantly associated with underground tissues (Fig. 4a, Supplementary Fig. S8; Supplementary Table S16). Among them, Y3-MEred simultaneously contained genes including *PnAACT1*, *PnFPPS*/*PnGGPS1*, *PnSS5* and *PnUGT18*, constituting the most complete coordinated unit supporting broad-spectrum ginsenoside accumulation in the main root and rhizome (Supplementary Table S16). Y3-MEred was significantly enriched in starch and sucrose metabolism and in amino sugar and nucleotide sugar metabolism (Fig. 4d, Supplementary Table S17), directly linking sugar donor generation to terminal saponin modification. In parallel, Y3-MEbrown, Y3-MEpink and Y3-MEsalmon were positively correlated with Rc, Rb2 and Rb3 and were more biased towards reproductive tissues. Y3-MEbrown was enriched in the TCA cycle, proteasome and ubiquitin-mediated proteolysis (Supplementary Table S17). It integrated genes including *PnHDS2*, *PnDXR*, *PnCYP9*/*37*/*76* and *PnUGT33*/*56* in flower tissues, and connected to transcription factors such as B3, BES1, bZIP, MYB, SBP and TCP (Fig. 4f).

To further dissect the internal drivers of these modules, we analyzed the expression patterns and functional annotations of the top 20 hub genes in each saponin related key module (Supplementary Fig. S9, Supplementary Table S18). These hub genes showed tissue-specific expression patterns that mirrored the tissue specificity of saponin accumulation. Notably, the hub genes of the key modules were annotated to include multiple transcription factors and functional proteins closely associated with ginsenoside accumulation and *UGT* mediated post-modification (Supplementary Table S18). Y1-MEblue and Y1-MEred in first year plants primarily contained basal regulatory nodes such as zinc finger MYM-type proteins and heat stress transcription factors (Fig. S9a,b). Y2-MEbrown, Y2-MEpink, and Y2-MEred in second year plants primarily contained transcription factors such as Dof zinc finger, SBP/SPL, CCCH/C2H2 zinc finger and bZIP, accompanied by important functional proteins including sugar transporters, MAPK, Exo70, and proteasome subunits (Fig. S9c,g). Y3-MEgreenyellow, Y3-MEpink, and Y3-MEblue in third year plants further exhibited regulatory and metabolic support nodes, including AP2/ERF-like, MYB DNA-binding, WAT1-related protein, BURP domain-containing protein, fibrillin, oxygen-evolving enhancer protein and STN7 kinase (Fig. S9h,l). Thus, the core of ginsenoside-related modules is not composed solely of structural genes, but is jointly supported by organ-specific transcriptional regulation, metabolite transport, signal transduction, and metabolic support systems. The genetic basis of ginsenoside accumulation presents as a spatiotemporally specific network that undergoes continuous rewiring with developmental progression: a foundational biosynthetic framework is established at the first year stage, whereas at the third year stage, modular specialization simultaneously enables broad-spectrum accumulation and branch-specific modification.

### Spatiotemporal expression differentiation of *UGTs* directly linked to differential ginsenoside accumulation profiles

From the key modules, we selected 17 differentially expressed *UGT* genes. These *UGTs* could be grouped into three spatiotemporal expression patterns: underground tissue-preferential (e.g., *PnUGT18*, *PnUGT60*), aerial tissue-preferential (e.g., *PnUGT33*, *PnUGT37*), and composite expression types (e.g., *PnUGT1*) (Fig. 5a). Among them, *PnUGT18* showed the highest expression in the rhizome, which increased with growth years. *PnUGT37* was stably expressed at high levels in leaf. And *PnUGT33* was enriched in leaf and flower, with its highest expression shifting to flowers in the third year.

**Fig. 5.**
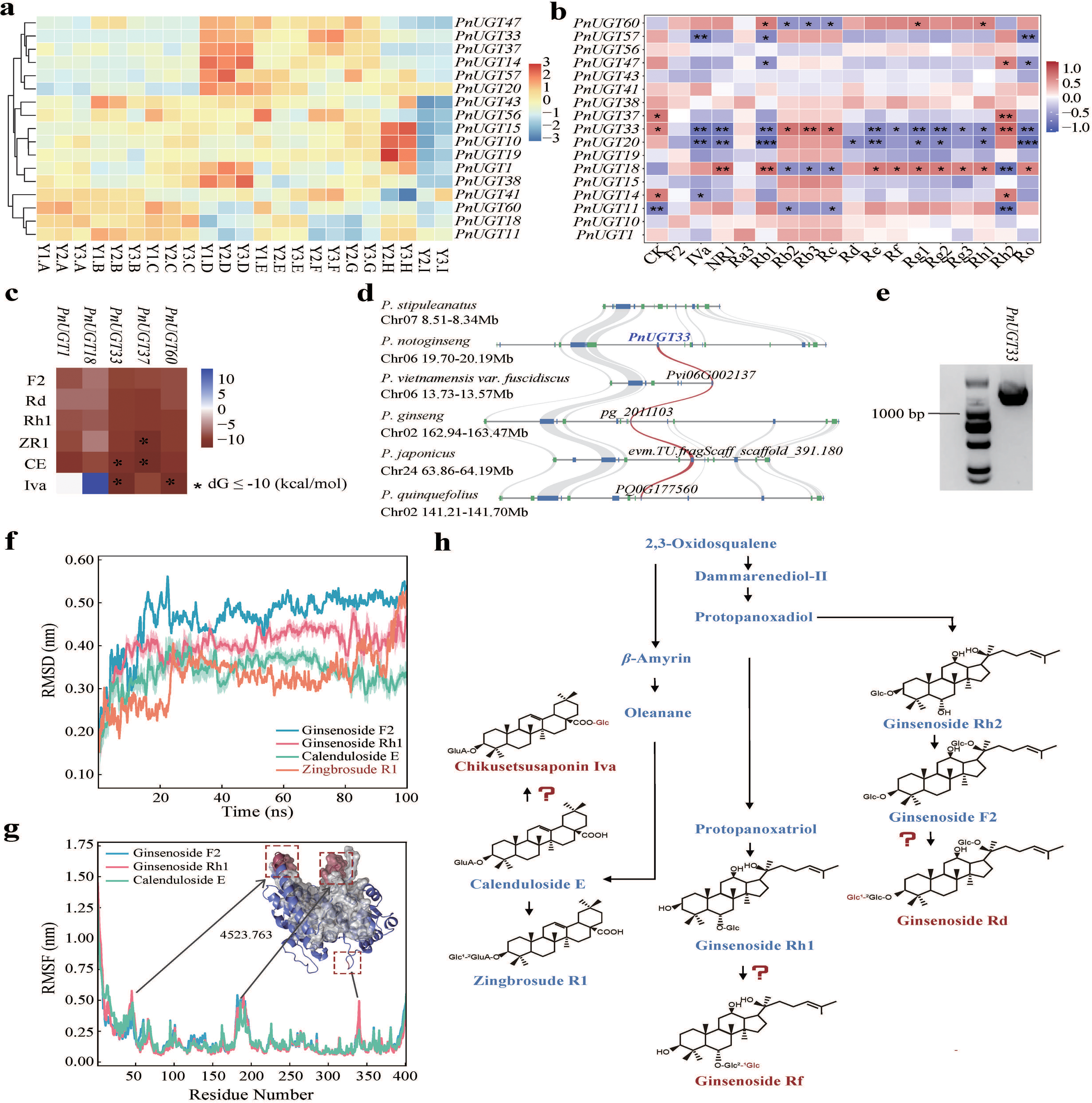
Characterisation of candidate *UGTs* involved in triterpene saponin biosynthesis. **(a)** Expression heatmap of *UGTs* in key modules. **(b)** Correlation analysis between key *UGTs* and saponin content. Colour scale indicates correlation strength; *p < 0.05, **p < 0.01. **(c)** Binding free energy of key candidate UGTs with substrates by molecular docking. Horizontal axis: different candidate UGTs; vertical axis: different saponin substrates. **(d)** Syntenic relationships of *PnUGT33* within *Panax* species. Collinear block shows the genomic region containing *PnUGT33* and homologous blocks in other *Panax* species, with chromosome and genomic locations annotated. **(e)** Cloning result of *PnUGT33*. The right lane shows the target gene band. **(f)** RMSD profiles of PnUGT33 in different molecular dynamics simulation systems. Unstable binding substrate is indicated in red. **(g)** RMSF distribution characteristics of PnUGT33 in different simulation systems. Regions with high fluctuations are marked by red boxes. **(h)** Chemical structures and potential glycosylation sites of substrates recognised by PnUGT33. Red question marks indicate potential glycosylation sites catalysed by PnUGT33.

Correlation analysis revealed clear correspondence between *UGT* expression and specific ginsenoside accumulation (58 out of 306 combinations reached FDR < 0.05). *PnUGT33* was positively correlated with most PPD-type ginsenosides (Rh2, CK, Rc, Rb2, Rb3) and negatively correlated with most PPT-type ginsenosides (Re, Rh1, Rg2, Rg1, Rf). *PnUGT18* showed the opposite pattern, being positively correlated with PPT-type ginsenosides (Rg1, Rg2, Re, Rh1, Rf) and some PPD-type ginsenosides (Rb1, Rg3), while negatively correlated with certain PPD-type ginsenosides (Rh2, Rb2, Rb3, Rc) (Fig. 5b, Table S19). *PnUGT20* was significantly negatively correlated with Rb1, Rd, Re, Rh1, Rg2, Rg1, Ro, IVa and NR1. *PnUGT60* was positively correlated with Rb1, Rg1, and Rh1, and negatively correlated with Rb2, Rc and Rb3. These results indicate that the terminal glycosylation mediated by different *UGT* members in *P. notoginseng* is closely associated with the differential accumulation profiles of distinct ginsenoside types.

Phylogenetic analysis revealed that these candidate *UGTs* originated from distinct evolutionary groups: *PnUGT37* was positioned within the clade related to *UGT1_Panax/PgUGT74AE2. PnUGT60* was within the clade related to *UGT73C33/UGT73F3/UGT73F24. PnUGT11* and *PnUGT33* belonged to the *UGT8* clade, and *PnUGT18* and *PnUGT20* were located in deep lineage-specific branches (Supplementary Fig. S4).

### PnUGT33 differentially recognises distinct triterpene skeletons, providing molecular evidence for UGT-driven metabolic branching

We selected the representative candidate gene *PnUGT33* to further evaluate its potential recognition capacity toward different ginsenoside substrates. Synteny analysis showed that the genomic region harboring *PnUGT33* was generally conserved across multiple *Panax* species (Fig. 5d). The gene contains a complete open reading frame of 1,206 bp, encoding 401 amino acids and exhibits the typical PSPG box of plant UGT enzymes (Fig. 5e, Tables S20 and S21). Molecular docking results showed that PnUGT33 exhibited favorable binding trends toward the oleanane-type saponins CE and IVa (−10.3 and −11.8 kcal/mol, respectively), while also maintaining relatively low binding energies toward the PPD-type F2 and Rd and the PPT-type Rh1 (−8.3, −8.9, and −8.8 kcal/mol, respectively) (Fig. 5c).

Based on these results, we selected four representative substrates F2, Rh1, CE and ZR1, for 100 ns molecular dynamics simulations. RMSD analysis showed that the PnUGT33-ZR1 complex system exhibited markedly elevated fluctuations during the later phase of the simulation, indicating relatively weak binding stability, whereas the complexes with the other three substrates maintained low and stable RMSD levels (Fig. 5f). RMSF analysis revealed that the overall backbone of PnUGT33 remained stable across the different systems; however, the F2 and Rh1 systems induced higher residue fluctuations in specific regions of the middle-to-latter segment of the protein (Fig. 5g), indicating that different substrates can differentially affect the local conformational flexibility of PnUGT33. MM/PBSA residue energy decomposition showed that favorable binding was primarily driven by van der Waals interactions. In the CE system, Ile127 and Tyr22 contributed most prominently (ΔG = −10.289 and −9.924 kJ/mol, respectively), while Phe25 served as a common contributing residue across all systems (Table S22), suggesting that these sites constitute an important structural basis for substrate recognition.

Taken together, these results demonstrate that PnUGT33 exhibits differential recognition capacity and binding stability toward ginsenosides of the PPD, PPT and oleanane backbone types. This further supports the notion that *UGT* members participate in the post-modification of distinct triterpenoid saponin branches through substrate selectivity, thereby contributing to the chemical diversification of ginsenosides in *P. notoginseng* (Fig. 5h).

## Discussion

Our study, integrating a chromosome-level genome assembly with spatiotemporal multi-omics analyses, systematically dissects the genetic basis of triterpene saponin diversity in *P. notoginseng*. The results show that the saponin diversity of *P. notoginseng* cannot be simply attributed to the expansion of core biosynthetic pathway gene families, but rather stems more likely from the lineage-specific differentiation of the UGT modifying enzyme family, the continuous rewiring of modular regulatory networks during development, and *UGT* mediated spatiotemporally specific post-modification processes.

### An evolutionary paradox: chemical diversity without gene family expansion

A prevailing model in plant specialized metabolism holds that gene duplication and family expansion provide the primary raw material for metabolic innovation^26^. In *Panax*, the telomere-to-telomere genome of P. ginseng identified 273 *UGTs* and showed that PD (Proximal duplication) and TD (tandem duplication) are major drivers of *UGT* duplication and divergence ^27^. Comparative genomics of 12 plant species further revealed that key enzyme genes in *Panax* species have followed independent evolutionary lineages, indicating unique functional adaptations^28^. In addition, systematic cloning and characterisation of multiple UGT94 family members from cDNA samples of *P. ginseng* revealed unprecedented sequence and functional diversity^29^. With their expanded gene families, tetraploid *Panax* species appear to conform to an expansion-driven model of metabolic evolution. *P. notoginseng*, however, presents a striking counterexample: despite harboring the largest number of species-specific gene families among the six *Panax* species examined, it shows no generalized expansion in most core pathway gene families; instead, multiple UGT subfamilies exhibit copy number contraction.

This evolutionary paradox suggests that ginsenoside diversification in diploid *P. notoginseng* may have followed an evolutionary trajectory distinct from that of the tetraploid species. Previous studies in tea plants have reported that the evolution and functional differentiation of glycosyltransferase genes have shaped tea quality and cold tolerance independently of whole-genome duplication^30^. Reconstruction of the evolutionary history of the plant UGT family has also revealed functional divergence of duplicated genes in specialised metabolism^31^. In *P. notoginseng*, the newly identified *UGTs* are dispersed across different lineages and lack syntenic relationships among subfamilies. This indicates that *UGT* evolution in this species tends to rely more on local duplication accompanied by sequence divergence and functional shift, rather than genome-wide tandem amplification. This strategy of fine-scale diversification may allow the diploid P. notoginseng to achieve a saponin structural diversity comparable to that of tetraploid species with a relatively compact *UGT* repertoire.

### Modular network rewiring and spatiotemporally compartmentalized ginsenoside accumulation

Previous transcriptomic studies of *P. notoginseng* provided initial clues about the tissue specificity of saponin biosynthesis. A comparative transcriptome analysis of leaves, roots and flowers identified candidate *CYP* and *UGT* genes involved in saponin biosynthesis in different tissues, and reported that the overall expression level of saponin biosynthetic genes was lower in flowers than in leaves and roots^5^. Full-length transcriptome sequencing further revealed tissue-specific transcript isoform distributions of genes involved in ginsenoside biosynthesis^32^. However, these studies are based on static comparisons of single tissues or single growth years and thus have not uncovered the dynamic logic underlying t he regulation of ginsenoside biosynthesis.

In our multi-year WGCNA data, we observed that the saponin biosynthesis network is not composed of a fixed set of co-expressed genes but is continuously rewired during development. In opium poppy (*Papaver somniferum*), WGCNA revealed spatiotemporal expression patterns and network rewiring of benzylisoquinoline alkaloid (BIA)-related genes at different developmental stages and in different tissues, indicating that metabolite diversification often depends on progressive modularisation of regulatory networks rather than linear regulation of a single conserved pathway^33^. In saffron, co-regulatory network analysis also revealed dynamic modular features of secondary metabolite biosynthesis during development^34^. The evolutionary trajectory observed in *P. notoginseng* in this study shifts from basic synthetic modules to mature patterns that integrate widespread accumulation and branch-specific modification. It is highly consistent with the above findings in concept. This indicates that development-dependent rewiring of co-expression networks may serve as a universal regulatory mechanism for the spatiotemporal diversification of plant secondary metabolism.

At the regulatory node level, we detected multiple transcription factors including AP2/ERF, MYB and bHLH in the key modules, which is consistent with a recent review proposing that bHLH and ERF transcription factors participate in the regulation of secondary metabolism in medicinal plants^35^. Moreover, hub gene analysis revealed non-enzyme proteins such as sugar transporters, MAPK signalling components and proteasome subunits, indicating that the saponin synthesis machinery is embedded in a broader cellular network encompassing transcriptional regulation, metabolite transport and protein turnover.

### Substrate promiscuity and branch selectivity as a distinctive strategy of UGTs in *P. notoginseng*

Substrate promiscuity of *Panax UGTs* has been widely reported. Using synthetic biology platforms, the complete pathways of major triterpene glycosylation products in *P. notoginseng* have been gradually resolved, showing that some *UGTs* have broad substrate adaptability^16^. RNA interference experiments have confirmed the functional regulatory role of *UGTs* in ginsenoside biosynthesis^15^. However, most previous studies focused on the breadth of the substrate spectrum of individual *UGTs*, with less attention paid to how substrate selectivity itself drives metabolic branching. A review proposed that enzyme promiscuity, functional mimicry and adaptive regulation are important drivers of the evolution of plant specialised metabolism^36^. Furthermore, it has been suggested that exploiting enzyme promiscuity to shape plant specialised metabolism is an important strategy in metabolic engineering^37^.

Our analysis of *PnUGT33* provides mechanistic clues as to how substrate promiscuity can be translated into branch selectivity. Although *PnUGT33* can accept substrates of the PPD, PPT and oleanane backbone types, its binding stability toward these substrates differs markedly: binding with the oleanane-type CE is the most stable, whereas the complex with ZR1 exhibits pronounced instability during the later phase of the simulation. This broad-spectrum yet non-equivalent recognition mode, which couples substrate promiscuity with intrinsic structural preference. It provides a direct biochemical basis for metabolic branching: within a given tissue context, the same UGT enzyme can preferentially glycosylate a specific substrate depending on the locally available precursor pool, thereby channeling metabolic flux toward a particular ginsenoside branch. In *Entada phaseoloides*, the substrate promiscuity of UGT73 family enzymes and their functional residues has also been shown to participate in similar metabolic branching regulation^38^. The prevalence of this mechanism suggests that the substrate selectivity of modifier enzymes, rather than mere substrate promiscuity, may be a key factor driving the structural diversification of plant specialised metabolism.

### An integrated model and future perspectives

Based on the above analyses, we propose a three-level integrated model to explain the chemical diversity of ginsenosides in *P. notoginseng*. At the **evolutionary level**, the UGT family expands its functional repertoire through lineage-specific differentiation rather than global expansion; recent LTR retrotransposon activity may have provided the structural basis for this local genomic plasticity by introducing new cis-regulatory elements or mediating chromosomal rearrangements. At the **regulatory level**, developmentally programmed modular rewiring of co-expression networks organises these *UGTs* into tissue- and stage-specific regulatory modules, establishing the spatiotemporal framework for differential saponin accumulation. At the **execution level**, differential expression of *UGTs* and their substrate recognition, which is both broad-spectrum and selective, translates the regulatory architecture into concrete metabolic branch outputs. A recent study in Sapindaceae provided direct evidence that LTR retrotransposons can drive the evolution and functional specialisation of triterpene saponin biosynthetic gene clusters^39^, indirectly supporting our inference that transposon activity participates in shaping the *UGT* regulatory network.

This model not only explains how the diploid *P. notoginseng* achieves high metabolic complexity with limited genomic resources, but also offers a new perspective on the fundamental difference in saponin diversification strategies between diploid and tetraploid *Panax* species. Tetraploids may rely more on gene copy number expansion to expand metabolic space, whereas diploids rely more on fine-scale innovation at the regulatory and modification levels. In the future, single-cell and spatial transcriptomics will help to resolve the precise spatial correspondence between *UGT* expression and saponin accumulation at cellular resolution, providing more accurate target genes for molecular breeding of medicinal plants.

## Methods

### Plant materials

A single individual of *P. notoginseng* used for genome sequencing was collected from Wenshan Prefecture, Yunnan Province, China. Fresh, healthy leaves were flash-frozen in liquid nitrogen and stored at −80 °C. Plants of first, second and third year *P. notoginseng* used for transcriptomic and metabolomic analyses were collected from the same region. Nine tissues – main root, fibrous root, rhizome, leaf, stem, flower, pedicel, berry and seed – were collected separately, immediately frozen in liquid nitrogen and stored at −80 °C. Because one-year-old plants are small and have not yet flowered, a pooled sampling strategy was used for this developmental stage. The samples used for metabolomic detection were exactly the same as those used for transcriptomic analysis.

### Genome sequencing and size estimation

High-quality genomic DNA was extracted using a modified CTAB method^40^. Paired-end libraries with an insert size of approximately 260 bp were constructed and sequenced on an Illumina HiSeq 2000 platform. In parallel, libraries with an insert size of approximately 20 kb were prepared according to the standard PacBio protocol and sequenced on a PacBio Sequel II platform. For Hi-C library construction, young leaves from the same individual were fixed with 1% (v/v) formaldehyde. Nuclei permeabilisation, chromatin digestion and proximity ligation were performed as described previously^41^. Hi-C libraries were finally sequenced on an Illumina HiSeq X Ten platform.

*K*-mer analysis was performed using quality-filtered Illumina reads. The frequency distribution of 17-mers was computed, and the genome size was estimated using GCE based on the formula: genome size = total number of *K*-mers / peak depth. GenomeScope was used to assess whole-genome heterozygosity^42^.

### Genome assembly and completeness assessment

The genome was assembled using a combination of Illumina short reads, PacBio long reads and Hi-C chromatin conformation capture data. De novo assembly was performed with Falcon^43^, using the longest 55× subreads as seeds for iterative correction (parameters: length_cutoff_pr = 7000, max_cov = 150). The resulting primary contigs (p-contigs) were polished sequentially: PacBio long reads were aligned to the contigs with pbalign and polished with Arrow; then Illumina short reads were aligned with BWA^44^ and two rounds of Pilon correction were performed^45^ to remove single-base errors and short indels. Residual allelic contigs were identified and reassigned using Purge Haplotigs^46^. Chromosome scaffolding was performed with the Lachesis pipeline using Hi-C data^47^. The Hi-C heatmap was visualised with Juicebox, and after manual curation the assembly was finalised into 12 high-confidence chromosomes^48^.

Genome completeness was assessed from multiple perspectives: (1) BUSCO was used to evaluate gene space completeness^49^; (2) high-quality Illumina reads were aligned to the assembly with BWA^44^; (3) previously obtained SMRT full-length transcripts were aligned to the assembly with GMAP^50^.

### Genome annotation

Protein-coding gene prediction integrated ab initio, homology-based and RNA-seq-based strategies. Using previously published full-length transcript data32 and approximately 30.9 Gb of clean RNA-seq data from root, stem, leaf, flower and rhizome, transcripts were assembled with Trinity^51^. Redundant transcripts with >90% similarity were removed with CD-HIT^52^, and gene structures were refined with PASA^53^. Manually curated gene models were used to train Augustus^54^and SNAP^55^. Ab initio predictions were performed on the hard-masked genome. Known protein sequences from several Apiales species (including *Coriandrum sativum*^56^, *Daucus carota*^57^, *P. ginseng*^58^ and *P. notoginseng*^2^) were aligned to the genome with Exonerate using the Protein2Genome model^59^. For RNA-seq-based prediction, transcripts were mapped to the soft-masked genome with GMAP^50^ and BLAT^60^ and iteratively refined by PASA. Finally, EVidenceModeler^52^ was used to integrate the evidence and generate a consensus gene set. Non-coding RNAs (rRNA, tRNA, snoRNA, snRNA and miRNA) were annotated with RNAmmer^61^, tRNAscan-SE^62^, snoScan^63^ and INFERNAL^64^, respectively.

### Repeat annotation

Repeat elements were identified using a combination of homology-based and ab initio searches. RepeatModeler was used for ab initio repeat family modelling. LTR_Finder^65^ and LTRharvest^66^ were run to identify LTR retrotransposons, and LTR_retriever^67^ was used to generate a high-quality LTR library. After merging the two libraries, repeats in the genome were annotated and masked with RepeatMasker (v4.0.5)^68^. Simple sequence repeats (SSRs) were detected with MISA using default parameters^69^.

For “full-length” LTR retrotransposons (those possessing both LTRs, a PBS and a PPT), open reading frames were classified using Pfam^70^ into Ty1-copia, Ty3-gypsy and unclassified groups. Amino acid sequences were aligned with ClustalW2^71^, sequences containing premature stop codons were removed, and a neighbour-joining tree was constructed with MEGA-X^72^.

### Analysis of comparative genomics

The *P. notoginseng* genome was compared with nine published genomes (*Daucus carota*^73^, *Centella asiatica*^74^, *Eleutherococcus senticosus*^75^, *Aralia elata*^76^, *P. vietnamensis* var. *fuscidiscus*^28^, *P. stipuleanatus*^77^, *P. ginseng*^27^, *P. quinquefolius*^77^ and *P. japonicus*^77^) using OrthoFinder (v2.5.2) to identify orthogroups^78^. A species tree was constructed using RAxML (v8.2.12) based on alignments of single-copy orthologous genes^80^, with the best substitution model selected by IQ-TREE (v2.2.3)^80^. Divergence times were estimated using MCMCTree in the PAML package using fourfold degenerate sites of single-copy genes (burnin=50,000, sampfreq=5, nsample=100,000). Gene family size dynamics were analysed with CAFE (v4.2.1)^81^. Functional annotation of genes was performed with EggNOG-mapper v2^82^; KEGG enrichment analysis was performed with clusterProfiler^83^. Synteny analysis was performed with the JCVI toolkit (Python version of MCscan).

### Transcriptome sequencing and differential expression analysis

Total RNA was extracted from each tissue sample using the RNAsimple Total RNA Extraction Kit (TIANGEN DP411). After quality assessment with a NanoDrop 2000 and a Bioanalyzer, paired-end sequencing was performed on an Illumina NovaSeq 6000 platform. Three independent biological replicates were used per tissue. Raw data were quality-controlled with FASTQC^84^ and adapters and low-quality reads (base quality ≤20 and proportion ≥27%) were filtered with fastp^85^. High-quality clean reads were aligned to our chromosome-level reference genome using HISAT2, and transcripts were assembled and quantified with StringTie^86^. Expression count matrices were obtained with HTseq^87^. Differential expression analysis was performed with DESeq2^89^ using |log₂ fold change| ≥ 1 and FDR ≤ 0.05 as thresholds.

After Z-score normalisation, genes with zero expression in all tissues were removed. Sample correlation analysis and K-means clustering were performed with the stats package in R; pathway enrichment of clustering results was performed with clusterProfiler^83^.

### Quantitative real-time RT-PCR validation

Total RNA was extracted using the PureLink™ Plant RNA Reagent (Thermo Fisher Scientific). Reverse transcription was performed using the FastKing RT Kit (TIANGEN, with gDNase). *PnActin2* was used as an internal control, and relative expression levels were calculated using the 2^(−ΔCt) method (three biological replicates and three technical replicates)^89^. PCR amplification was performed on a QuantStudio 5 real-time PCR system (Thermo Fisher) using TB Green® Premix Ex Taq™ II (Takara) Primers are listed in Supplementary Table S14.

### Identification of transcription factors and saponin biosynthetic genes

Using the Pfam database of transcription factors from PlantTFDB (https://planttfdb.gao-lab.org/) together with Table S8 as references, we identified triterpene saponin biosynthetic genes and transcription factors (with an E-value threshold of 1e-15) and visualized the resulting co-expression network using Cytoscape (v3.9.1). To construct a comprehensive phylogenetic tree, we integrated UGT sequences from the above-mentioned ten species along with data from our collected UGT dataset (Table S11). A maximum-likelihood tree was built using IQ-TREE^80^, which automatically selected the best evolutionary model and performed 1,000 bootstrap replicates. The tree was annotated using iTOL^90^. Subsequently, protein sequences of previously reported ginsenoside biosynthesis-related genes in P. notoginseng were downloaded from the NCBI database (Table S15). Using BLAST, we screened for homologs with thresholds of E-value ≤ 1e⁻¹⁰, query coverage ≥ 50%, and sequence identity ≥ 50%. The expression profiles of the identified homologs were visualized using the pheatmap package (1.0.13) in R.

### Metabolite extraction, detection and data analysis

Targeted metabolomics was performed on the same samples used for transcriptomics. Extraction and analysis were performed following a published method^91^ with minor adjustments. Briefly, tissue powder was ground in liquid nitrogen, and 0.05 g of sample was suspended in 10 mL of 6 mol/L HCl, mixed and frozen under vacuum. Hydrolysates were incubated at 110 °C for 22 h, cooled to room temperature, filtered, and 0.2 mL of the filtrate was evaporated at 60 °C, redissolved in amino acid buffer and subjected to LC-MS analysis.

Missing values in the metabolomic data were filled with zeros. Data visualisation was performed with the pheatmap package. Sample correlations were analysed using Pearson’s correlation coefficient. Principal component analysis (PCA) of saponin components and correlation analysis among components were performed with the R stats package.

### WGCNA and candidate UGT selection

Weighted gene co-expression networks were constructed separately for first, second and third plants using the WGCNA package^92^. After filtering low-variance genes, parameters were set as maxBlockSize=40,000, mergeCutHeight=0.15 and minModuleSize=20; a soft threshold (R²≥0.85) was chosen to make the network approximate a scale-free distribution. Module eigengenes were used to calculate correlations between modules and traits (saponin content and tissue type). Functional enrichment of module genes was performed with clusterProfiler^83^.

Candidate *UGTs* were selected according to two stringent criteria: (1) intersection of differentially expressed *UGT* genes in the significant module associated with the trait; (2) evaluation of Spearman correlations between *UGT* expression levels and saponin contents. Original data were log₂(x+1) transformed, and each gene–metabolite pair was tested for correlation; multiple testing was corrected using the Benjamini–Hochberg method (FDR). An FDR < 0.05 was considered significant, and |ρ| ≥ 0.5 was used as a threshold for strong correlation. Results were visualised with custom heatmaps. All analyses were performed in R (v4.3.2).

### Cloning of candidate *UGT* genes

PCR was performed using TIANGEN Taq DNA polymerase with cDNA as template. Primer information is given in Supplementary Table S20. PCR products were separated by 1% agarose gel electrophoresis, excised and purified (Steady Pure Agarose Gel DNA Purification Kit, Accurate Biology). Purified products were cloned into the pMD-19T vector (Takara) and transformed into DH5α chemically competent cells (Weidi). Recombinant plasmids were verified by colony PCR and then Sanger-sequenced by BGI to confirm the target sequences.

### Molecular docking and molecular dynamics simulations

Protein three-dimensional structures were built using AlphaFold2 (v2.3.2). Initial geometries of saponin substrates were downloaded from the PubChem database (https://pubchem.ncbi.nlm.nih.gov/). Molecular docking was performed with AutoDock Vina (v1.1.2)^93,94^; the docking pose with the best score and a reasonable conformation was selected as the starting coordinate for molecular dynamics simulations.

Molecular dynamics simulations were performed using GROMACS 2022^95^ with the CHARMM36 force field. Each protein–ligand complex was placed in a dodecahedral periodic box with a distance of at least 1.0 nm from the solute to the box edge. The system was solvated with TIP3P_CHARMM water molecules, and Na⁺ or Cl⁻ ions were added to neutralise the net charge. Hydrogen bond lengths were constrained using the LINCS algorithm^96^. Each system was first energy-minimised, then pre-equilibrated under NVT (100 ps) and NPT (100 ps) ensembles. Finally, a 100 ns production simulation was performed under the NPT ensemble with an integration time step of 2 fs. Simulation trajectories were analysed with GROMACS built-in tools and visualised with DuIvyTools (v0.4.8)^97^.

## Data availability

Genome data have been deposited in the NCBI BioProject database under accession number **PRJNA1427891**. RNA-Seq data have been deposited under **PRJNA1314388**. Targeted metabolomic data have been submitted to the NGDC under accession number **PRJCA058824**.

## Code availability

Custom scripts used for data analysis are available from the corresponding author upon reasonable request.

## Supporting information

Supplementary Large Tables

## Acknowledgements

This work was supported by the Natural Science Foundation of China (U1902205) and a startup grant of Hainan University to Li-zhi Gao. We would thank Wei Li and Ju-jin Jia for their technical assistance.

## Author contributions

L.-z. G. conceived the study. Z.X. and W.L. performed the experiments and analyzed data. F.-g. W., G. X. and Z.-j. C. collected the materials. Z.X. and L.-z. G. drafted the manuscript. L.-z. G. revised the manuscript. All authors read and approved the final manuscript.

