## Supplementary Figures and Tables for "A Diploid *Panax* Genome Reveals Ginsenoside Diversity Driven by UGT Family Diversification and Network Rewiring Rather than Gene Family Expansion": Supplementary Figures and Tables.docx

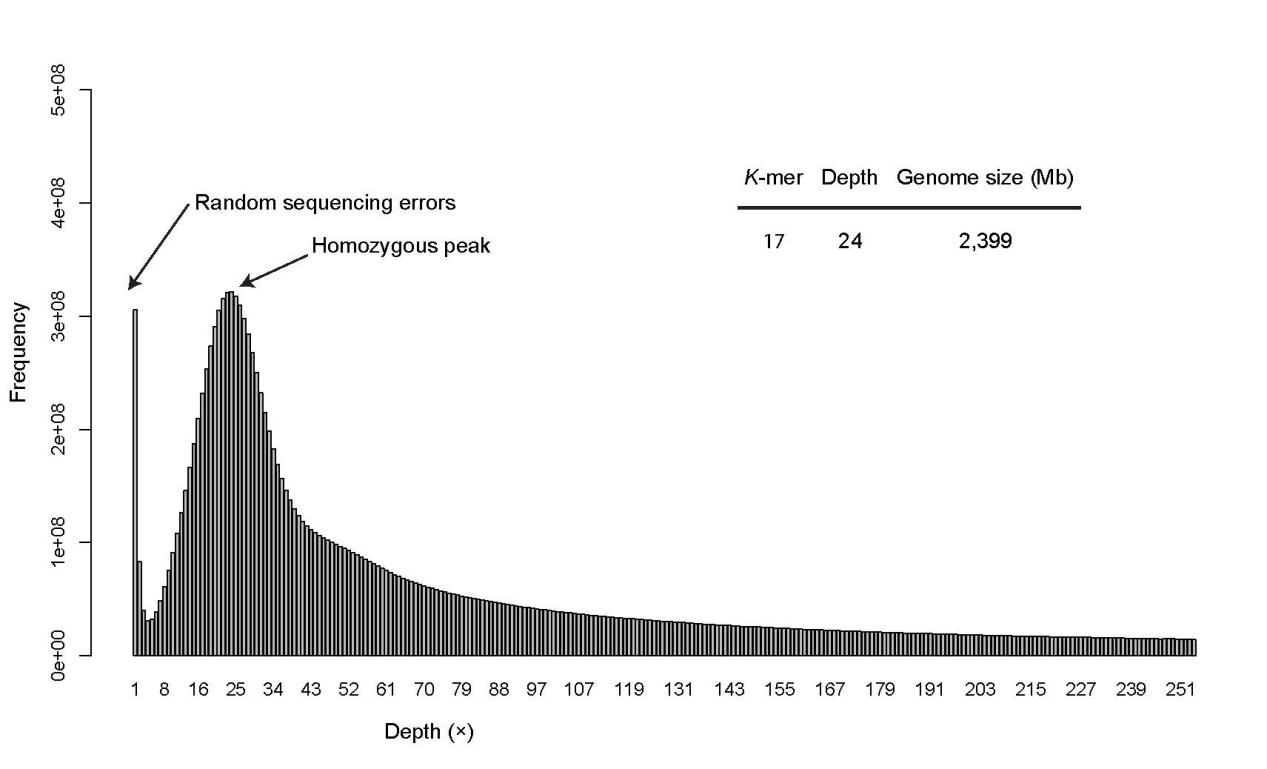

**Figure S1. The 17-mer distribution of sequencing reads from the *Panax notoginseng* genome**.

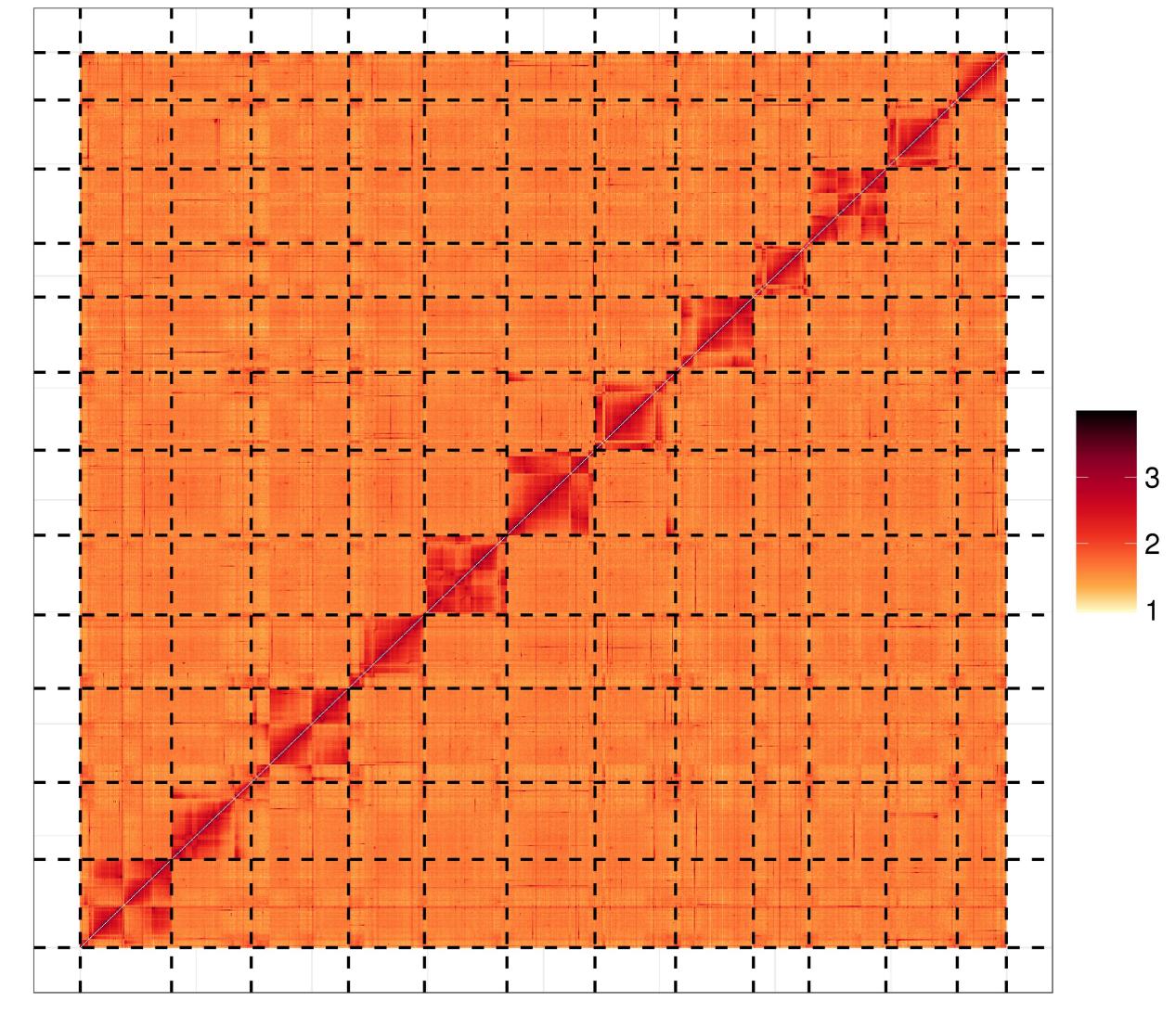

**Figure S2. Genome-wide all-by-all Hi-C interaction of *P. notoginseng*.**

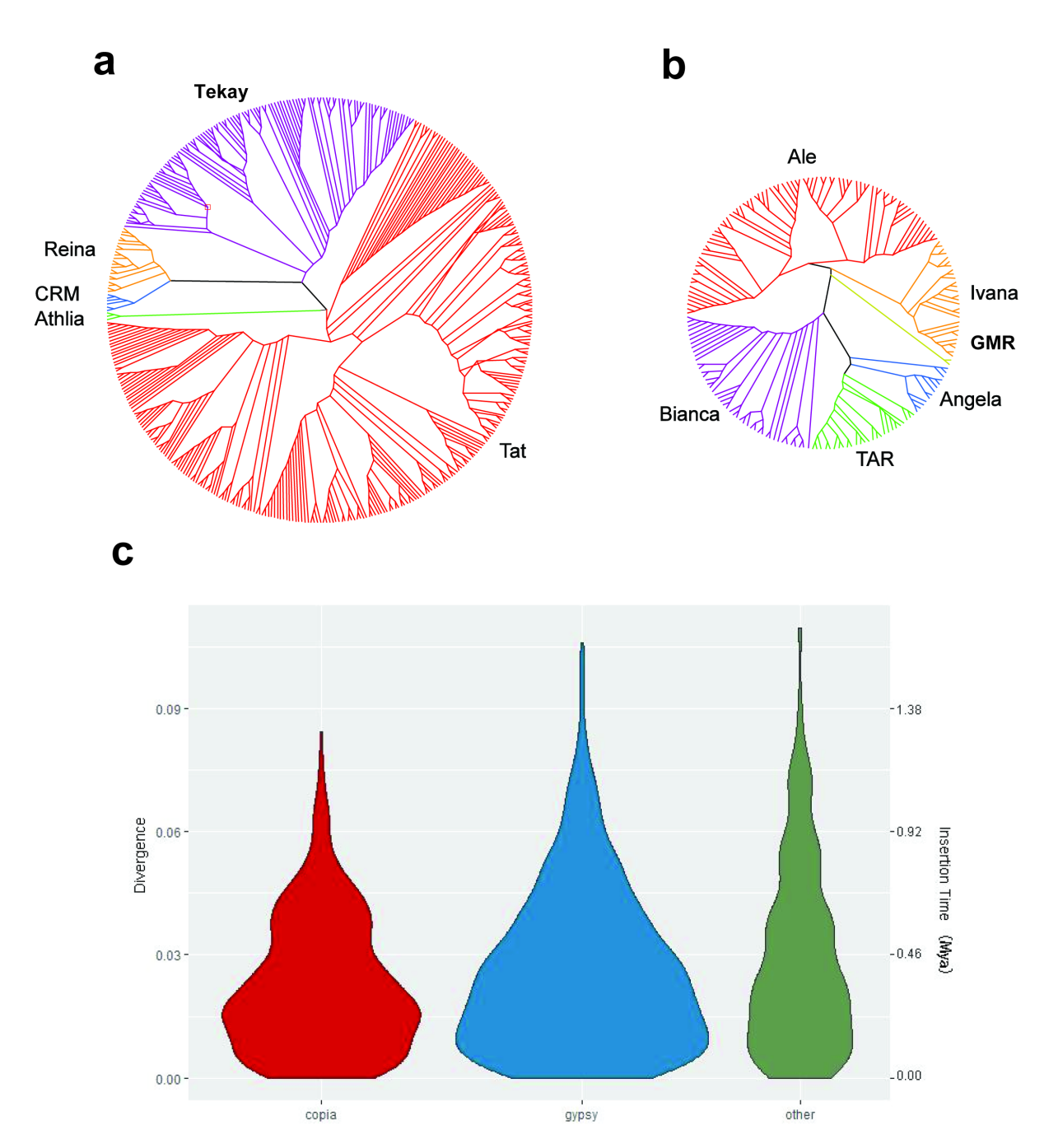

**Figure S3. Landscape of TEs in *P. notoginseng* genome.** The neighbor-joining and unrooted phylogenetic trees were constructed on the basis of 446 intact Ty3-*gypsy* **(a)** and 194 intact Ty1-*copia* **(b)** aligned sequences corresponding to the RT domains without a premature termination codon **(c)** The LTRs divergence of intac LTR-RTs in *P. notoginseng* genome.

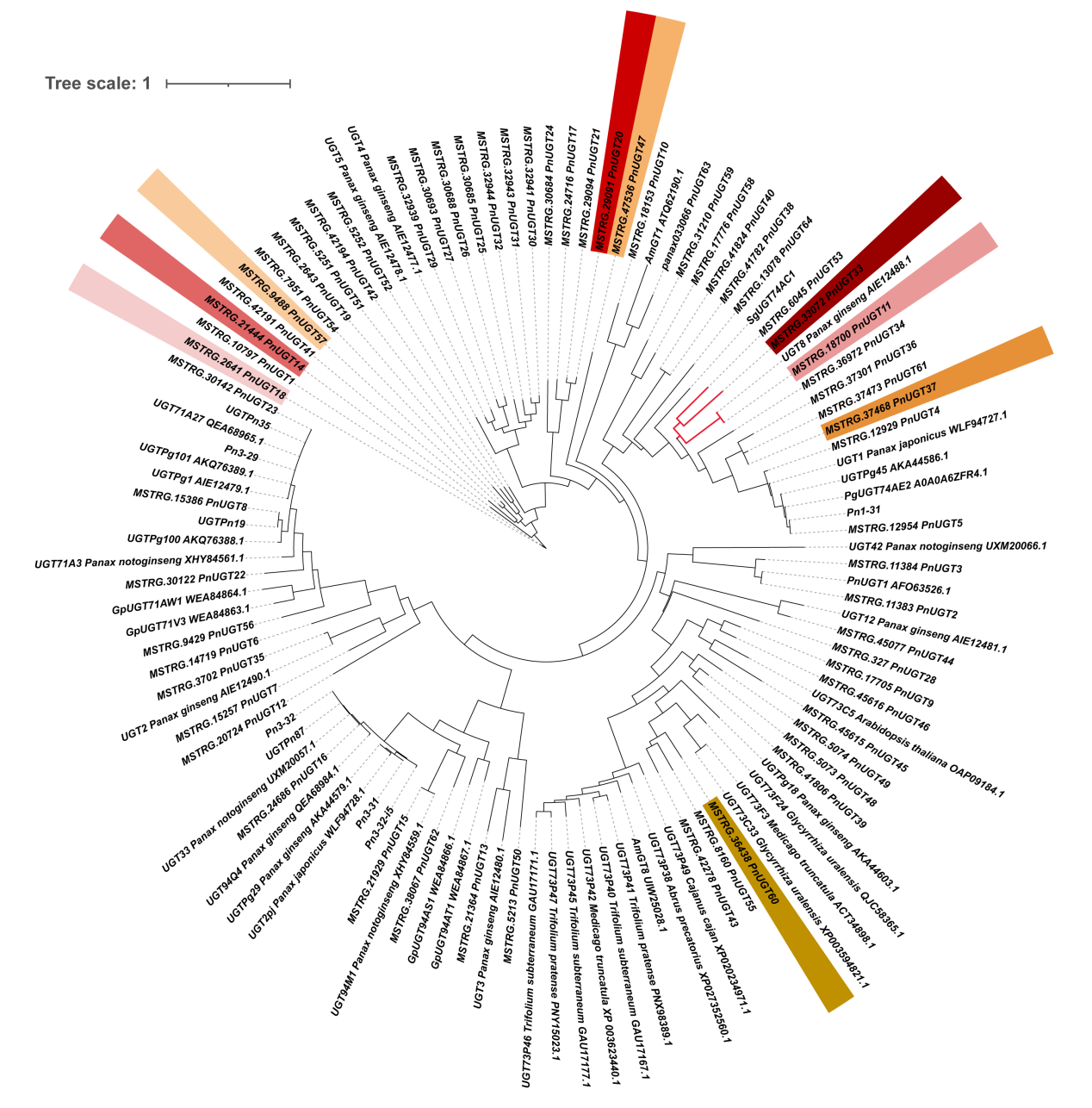

**Figure S4. Phylogenetic tree of UGT genes from *P. notoginseng* and other *Panax* species**

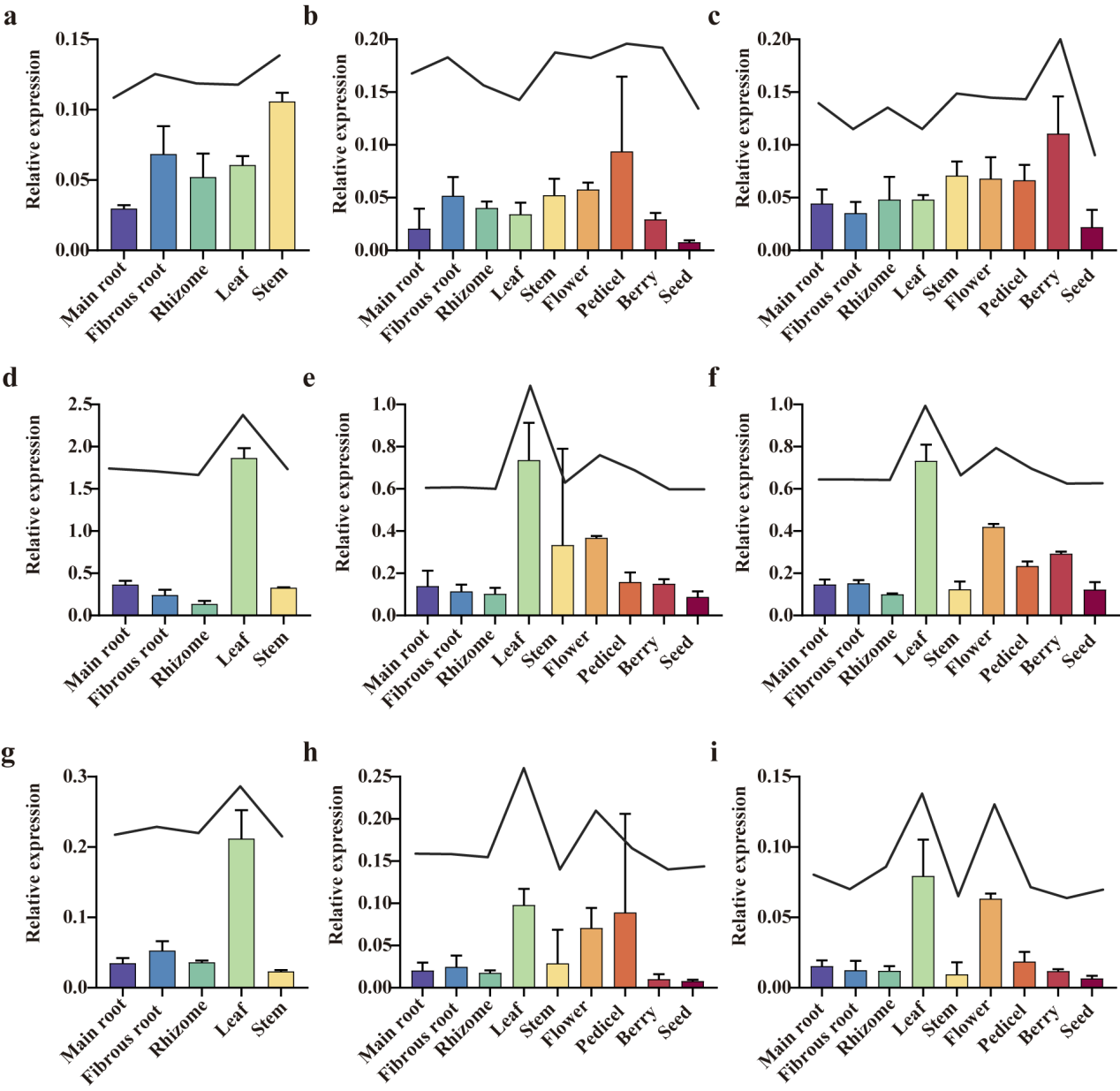

**Figure S5. qRT-PCR identification of transcriptome sequencing. (a)** PnUGT22 (Year 1); **(b)** PnUGT22 (Year 2); **(c)** PnUGT22 (Year 3); **(d)** PnUGT37 (Year 1); **(e)** PnUGT37 (Year 2); **(f)** PnUGT37 (Year 3); **(g)** PnUGT62 (Year 1); **(h)** PnUGT62 (Year 2); **(i)** PnUGT62 (Year 3).

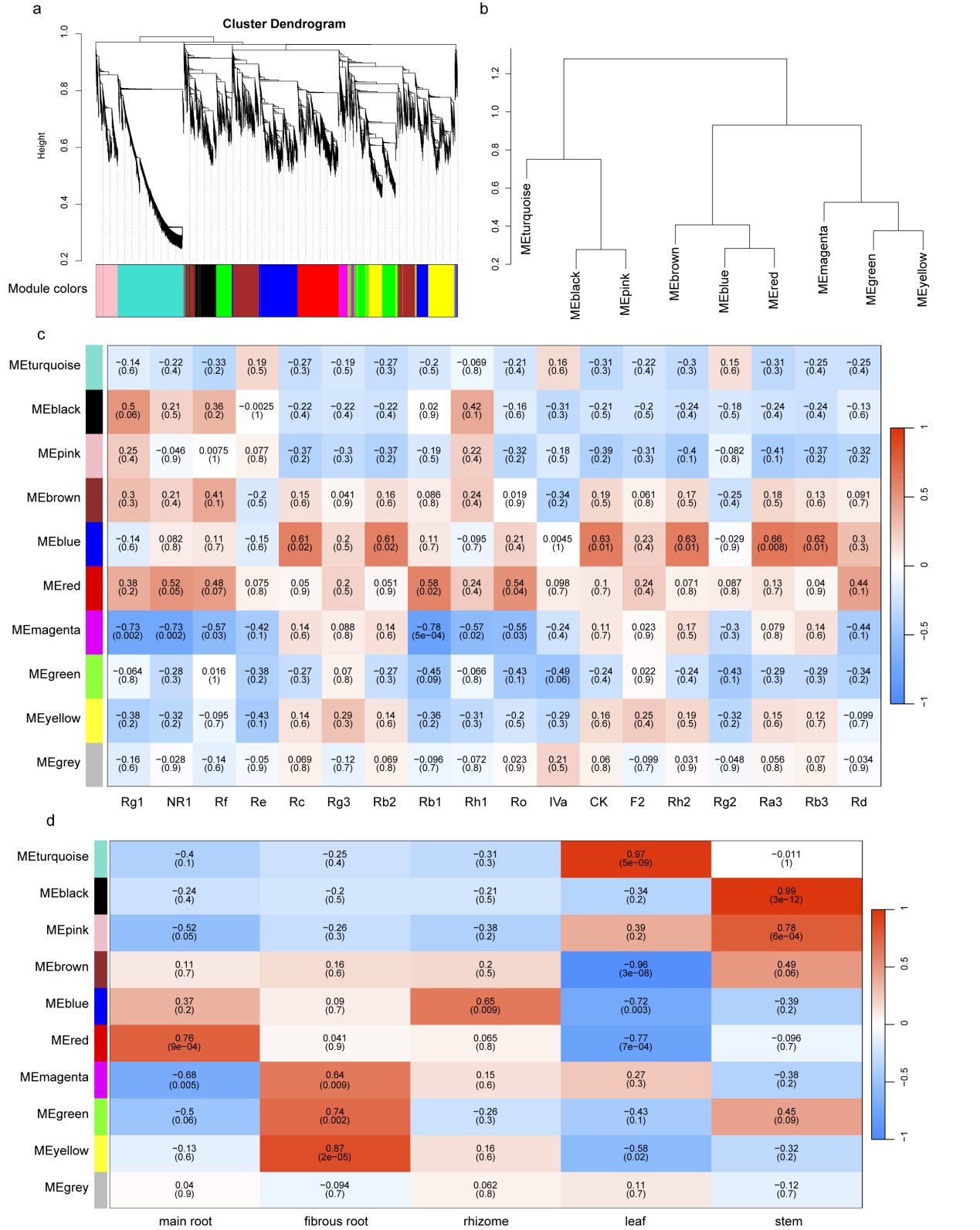

**Figure S6. Weighted gene co-expression network analysis (WGCNA) of *P. notoginseng* (Year 1). (a)** Hierarchical cluster tree showing coexpression modules identified by WGCNA; **(b)** Clustering dendrogram of module eigengenes; **(c)** Module-saponin content association; **(d)** Module-tissue association.

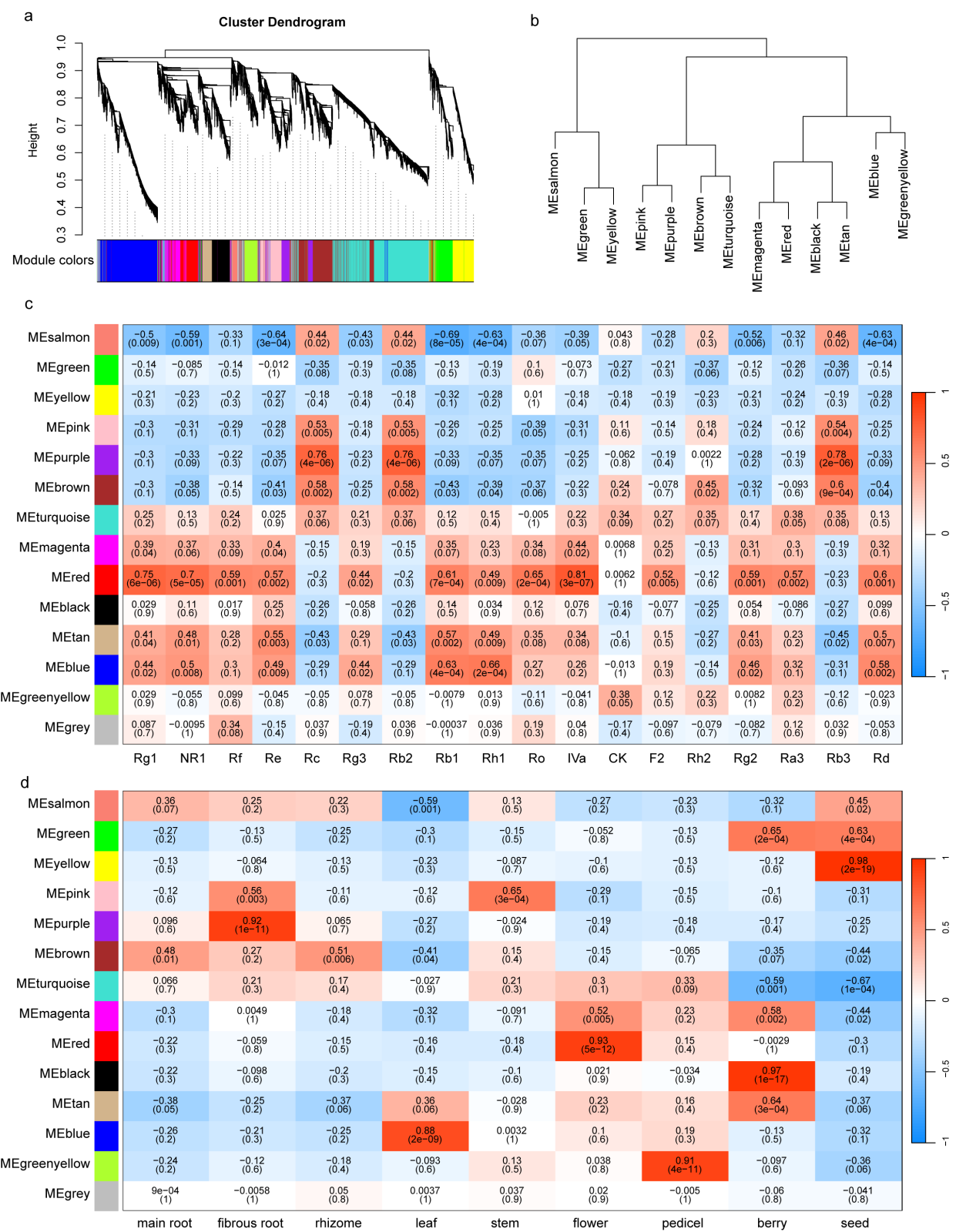

**Figure S7. Weighted gene co-expression network analysis (WGCNA) of *P. notoginseng* (Year 2). (a)** Hierarchical cluster tree showing coexpression modules identified by WGCNA; **(b)** Clustering dendrogram of module eigengenes; **(c)** Module-saponin content association; **(d)** Module-tissue association.

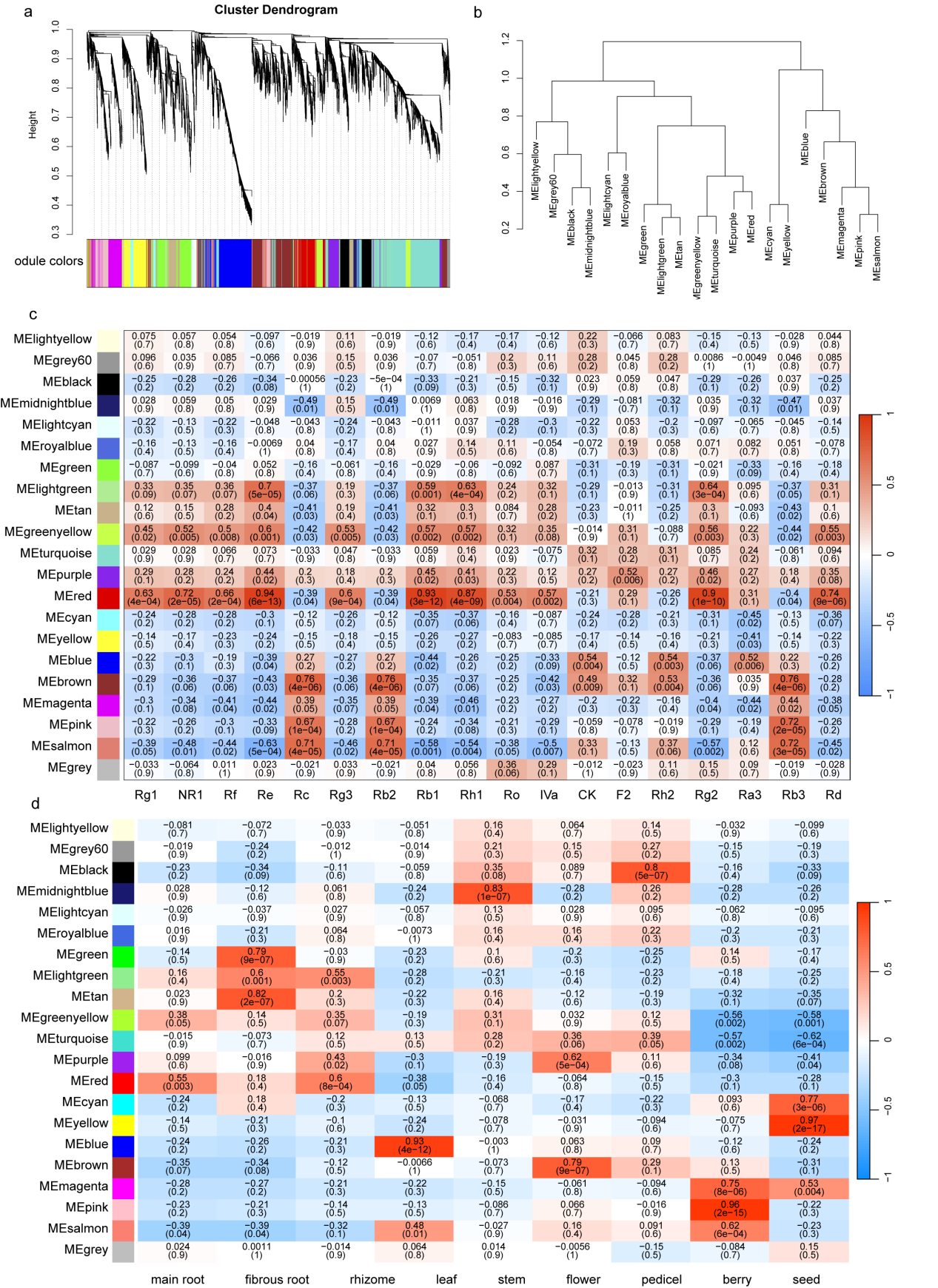

**Figure S8. Weighted gene co-expression network analysis (WGCNA) of *P. notoginseng* (Year 3). (a)** Hierarchical cluster tree showing coexpression modules identified by WGCNA; **(b)** Clustering dendrogram of module eigengenes; **(c)** Module-saponin content association; **(d)** Module-tissue association.

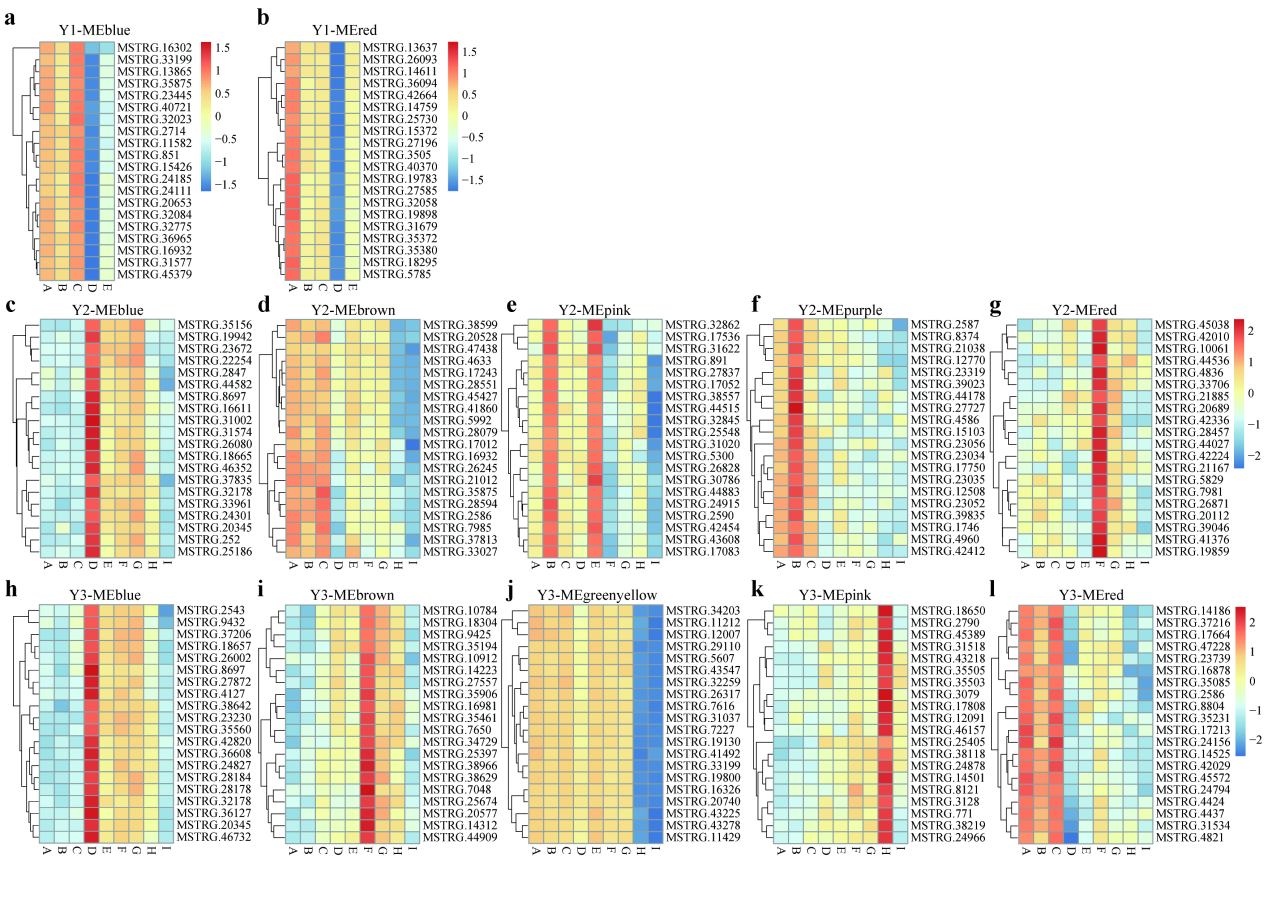

**Figure S9. Expression heatmap of hubd genes in WGCNA-related module pathways. (a-b)** Year 1; **(c-g)** Year 2; **(h-i)** Year 3.

**Table S1. Libraries and read statistics used for the *Panax notoginseng* genome assembly*.**

| **Sequencing technique** | **Insert Size (bp)** | **Average Read Length (bp)** | **Raw Data (Gb)** | **Raw Sequence Coverage (×)** |
| --- | --- | --- | --- | --- |
| NGS | 260 | 150 | 326.25 | 135.99 |
| Pacbio | 40,000 | 10,133 | 203.00 | 84.62 |

*The estimated genome size is ~2,399 Mb.

**Table S2. Assembly statistics of the *P. notoginseng* genome.**

| **Features** | **p-contigs** | **pseudo-chromosomes** |
| --- | --- | --- |
| Genome size (bp) | 2,270,879,957 | 2,271,278,847 |
| Contig number | 4,175 | 4,249 |
| Max contig length (bp) | 14,201,109 | 14,201,109 |
| Contig N50 (bp) | 1,109,761 | 1,083,164 |
| Contig N90 (bp) | 280,691 | 277,043 |
| Scaffold number | 4,175 | 259 |
| Max scaffold length (bp) | 14,201,109 | 219,072,531 |
| Scaffold N50 (bp) | 1,109,761 | 199,803,215 |
| Scaffold N90 (bp) | 280,691 | 153,439,021 |
| Gap number | 0 | 3,990 |
| Gap length (bp) | 0 | 399,000 |
| GC content (%) | 34.38 | 34.38 |

**Table S3. Chromosome lengths of the assembled *P. notoginseng* genome.**

| **Chromosome ID** | **Contig number** | **Chromosome length (bp)** |
| --- | --- | --- |
| 1 | 416 | 217,575,036 |
| 2 | 345 | 218,222,556 |
| 3 | 375 | 219,072,531 |
| 4 | 353 | 199,803,215 |
| 5 | 336 | 203,826,934 |
| 6 | 325 | 207,529,616 |
| 7 | 270 | 159,224,650 |
| 8 | 329 | 167,423,711 |
| 9 | 320 | 179,198,802 |
| 10 | 343 | 160,807,860 |
| 11 | 351 | 147,349,750 |
| 12 | 226 | 153,439,021 |
| Unanchored | 260 | 37,805,165 |
| Total | 4,249 | 2,271,278,847 |

**Table S4. Assessment of the *P. notoginseng* genome assembly using BUSCO.**

| **BUSCOs from Embryophyta lineage*** | | |
| --- | --- | --- |
| **Type** | **Number** | **Percent (%)** |
| Complete | 1,326 | 92.1 |
| Duplicated | 137 | 9.5 |
| Fragmented | 29 | 2.0 |
| Missing | 85 | 5.9 |

* The 1,440 BUSCO conserved genes used were collected from Embryophyta lineage;

### **Table S5. Validation of the *P. notoginseng* genome using Illumina and full-length transcripts.**

|  | **Total** | **Mapped** | **Rate (%)** |
| --- | --- | --- | --- |
| **Reads mapping** | | | |
| Short libraries | 730,788,014 | 724,365,899 | 99.12 |
| **Full-length transcripts*** | | | |
| Transcripts | 51,040 | 43,633 | 85.49 |

* Hits with Coverage >= 90% and Identity >= 90%

**Table S6. Summary of gene prediction for *P. notoginseng.***

| **Type** |  | **Values** |
| --- | --- | --- |
| **Gene Features** | Total number of predicted genes (#) | 46,425 |
|  | Average gene length (bp) | 3,793 |
|  | Average CDS length (bp) | 1,071 |
|  | Average exon per gene | 4.8 |
|  | Average exon length (bp) | 223 |
|  | Average intron length (bp) | 717 |
|  | tRNAs | 717 |
|  | rRNAs | 431 |
|  | snoRNAs | 108 |
|  | snRNAs | 407 |
|  | miRNAs | 137 |
|  | Transposable elements (%) | 87.51 |

**Table S7. Comparisons of repetitive sequence categories and contents among the *P. notoginseng* genomes.**

|  |  | **Fregments No.** | **Length (bp)** | **Percentage (%)** |
| --- | --- | --- | --- | --- |
| **DNA** | CMC-EnSpm | 30518 | 21386241 | 0.94 |
|  | hAT | 29682 | 9937521 | 0.44 |
|  | MULE-MuDR | 16623 | 9363299 | 0.41 |
|  | PIF-Harbinger | 3156 | 1420137 | 0.06 |
|  | TcMar-Stowaway | 20672 | 4222089 | 0.19 |
|  | Other | 7643 | 9267369 | 0.41 |
| **RC** | RC/Helitron | 5517 | 4057033 | 0.18 |
| **RNA** | LINE | 15129 | 15460880 | 0.68 |
|  | SINE | 153 | 14870 | 0.00 |
|  | LTR-Copia | 200786 | 185554456 | 8.17 |
|  | LTR-Gypsy | 645234 | 1367819705 | 60.22 |
|  | LTR-Other | 75442 | 45947869 | 2.02 |
| **Other Repeats** | Other_Repeats | 626118 | 293759311 | 12.93 |
|  | Simple_repeat | 358194 | 17840168 | 0.79 |
|  | Low_complexity | 30038 | 1574715 | 0.07 |
| **Total** |  | 2064905 | 1987625663 | 87.51 |

**Table S9. Summary of the gene family clustering.**

| **Species** | **Gene number** | **Unclustered gene number** | **Family number** | **Unique families** |
| --- | --- | --- | --- | --- |
| *P. stipuleanatus* | 39043 | 3093 | 21269 | 555 |
| *P. notoginseng* | 58369 | 11703 | 19373 | 3637 |
| *P. vietnamensis* var. *fuscidiscus* | 36454 | 2198 | 20869 | 231 |
| *P. japonicus* | 72503 | 8976 | 26767 | 1336 |
| *P. ginseng* | 77266 | 6674 | 27114 | 843 |
| *P. quinquefolius* | 64247 | 6880 | 25400 | 903 |

**Table S14. Sequences of primers used in the qRT-PCR analysis**

| *Gene name* | *Forward primer (5’-3’)* | *Reverse primer (5’-3’)* |
| --- | --- | --- |
| *PnActin2* | *TCCAAGGGTGAATATGATGAATCG* | *AACCTCTCCAAAGAGAATTTCTGAGT* |
| *PnUGT22* | *CCGCACTTCAGGTTTTTAGAAGTACC* | *CTCTCACGTGTGCCTTTTGGG* |
| *PnUGT37* | *GGCCAATTAAGACCATTGGACC* | *TGCATTGAGCCACTTGATGCAAG* |
| *PnUGT62* | *GGCCTACCACACCATCTTCAC* | *GATGCTATTTCCGGTACCCATGGC* |
